# Exploring functional protein covariation across single cells using nPOP

**DOI:** 10.1101/2021.04.24.441211

**Authors:** Andrew Leduc, R. Gray Huffman, Joshua Cantlon, Saad Khan, Nikolai Slavov

## Abstract

Many biological processes, such as the cell division cycle, are reflected in protein covariation across single cells. This covariation can be quantified and interpreted by single-cell mass-spectrometry (MS) with sufficiently high throughput and accuracy. Towards this goal, we developed nPOP, a method that uses piezo acoustic dispensing to isolate individual cells in 300 picoliter volumes and performs all subsequent sample preparation steps in small droplets on a fluorocarbon-coated slide. This design enabled simultaneous sample preparation of thousands of single cells, including lysing, digesting, and labeling individual cells in volumes of 8-20 nl. Protein covariation analysis identified cell-cycle dynamics that were similar across cell types and dynamics that differed between cell types, even within sub-populations of melanoma cells defined by markers for drug-resistance priming. The melanoma cells expressing these markers accumulated in the G1 phase of the cell cycle, displayed distinct protein covariation across the cell cycle, accumulated glycogen, and had lower abundance of glycolytic enzymes. The non-primed melanoma cells exhibited gradients of protein abundance and covariation, suggesting transition states. These results were validated by different MS methods. Together, they demonstrate that protein covariation across single cells may reveal functionally concerted biological differences between closely related cell states.

## Introduction

Single-cell measurements are commonly used to identify different cell types from tissues composed of diverse cells^1, 2^. This analysis is powering the construction of cell atlases, which can pinpoint cell types affected by various physiological processes. This cell classification requires analysing large number of cells and may tolerate measurement errors^1, 3, 4^.

In addition to classifying cells by type, single-cell measurements may reveal regulatory processes within a cell type and even associate them with different functional outcomes^5–7^. For example, the covariation among proteins across single cells from the same type may reflect cell intrinsic dynamics, such as the cell division cycle^5, 8, 9^. Furthermore, protein covariation may reflect protein interactions within complexes or cellular states, such as senescence and drug resistance^5^. However, estimating and interpreting protein covariation within a cell type requires high quantitative accuracy and high throughput^5, 10^. Indeed, protein differences within a cell type are smaller than across cell types and can be easily swamped by batch effects and measurement noise.

Therefore, we sought to minimize measurement noise to levels consistent with estimating and interpreting protein covariation across single cells from the same cell type. Towards this goal, we aimed to increase the number of cells prepared in parallel and thus reduce the number of batches and associated background noise that undermine the accuracy of single-cell proteomics by mass spectrometry (MS)^11–15^. Specifically, we aimed to develop a widely accessible, inexpensive, robust, and automated sample preparation method that reduces volumes to a few nanoliters.

Multiple methods for single-cell sample preparation have been developed^16–20^, but the use of tubes and wells has limited the number of samples that can be simultaneously prepared. Thus, our goal was to perform parallel sample preparation of thousands of single cells to increase the size of experimental batches^12, 21, 22^. To achieve this, we abandoned the traditional use of multi-well plates or chips and introduced a new design, droplets on a flat surface. This design allows for high density of samples with minimized volumes. It also avoids movement of the samples during sample preparation so that small volumes of reagents can be repeatedly and precisely dispensed to each droplet containing a single cell. These design considerations of nPOP are general and may be implemented by different liquid handlers. Here, we demonstrate them using the CellenONE cell sorting and liquid handling system.

nPOP enables preparing over 2,000 single-cell samples in one batch. Using this increased throughput, we study protein covariation within a cell type, and even within sub-populations of melanoma cells. The protein covariation identified biological processes that covary with the cell division cycle (CDC) in a cell type dependent and independent manner. Furthermore, this analysis identified melanoma clusters that differed in the abundance of markers for drug resistance. These clusters differed in metabolism and cell cycle dynamics. Proteins found to be differential between these two clusters showed a gradient of protein abundances within a cluster, suggesting a gradual transition to drug-resistance primed state that involves proteins functioning in metabolism and energy production.

## Results

### nPOP enables flexible droplet sample preparation

To increase throughput and reduce background noise, our goal was to maximize the number of single cells prepared in parallel while minimizing the reaction volumes used in sample preparation. To this end, we explored the idea of performing all sample preparation steps in droplets on the surface of a uniform glass slide, Fig. 1a and Extended Data Fig. 1a-c. Obviating wells used by other methods, nPOP achieves higher spatial density of samples (504 per glass slide using AL-01) and thus higher throughput (2,016 samples per preparation using AL-01). The droplet layouts also allow the freedom to arrange single cells in any geometry that best fits the experimental design, Fig. 1a,b and Extended Data Fig. 2. Supplemental software 2 provides 4 predefined layouts optimized for different experimental designs, including label-free MS^23^, SCoPE2^14^, and plexDIA^24^. Additional layouts can be easily defined for customized workflows.

**Figure 1.**
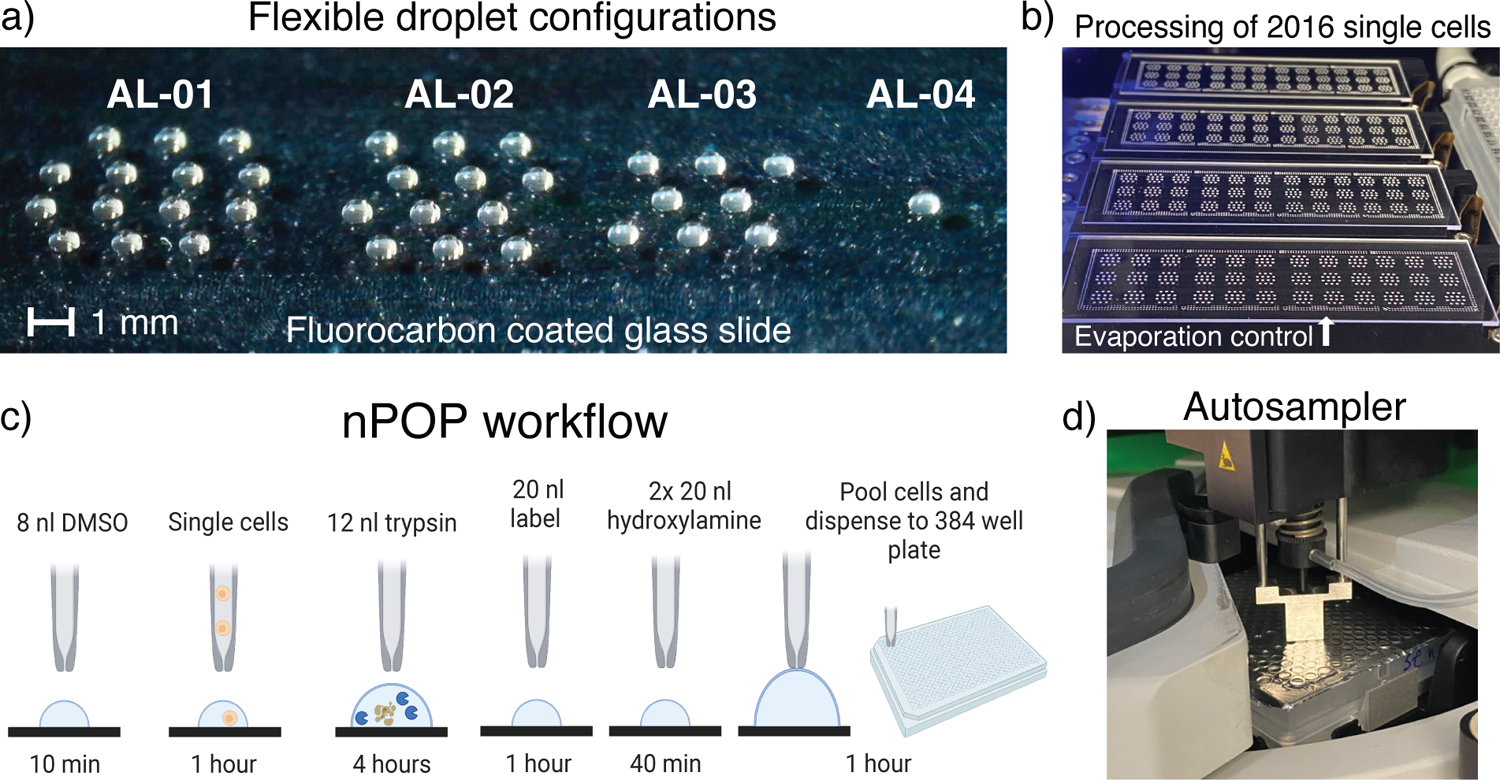
Parallel preparation of thousands of single cells by nPOP. (**a**) The slide based approach allows programming different droplet layouts, such as the 4 examples shown in the picture. (**b**) Four slides with 2,016 single cells from an nPOP experiment using droplet configuration AL-01. Samples are surrounded by a perimeter of water droplets for local humidity control. Slides are placed on a cooling surface to further prevent evaporation. (**c**) A schematic of the nPOP method illustrates the steps of cell lysis, protein digestion, peptide labeling with TMTpro, quenching of labeling reaction, and sample collection. These steps are performed for each single cell (corresponding to a single droplet). (**d**) After labeling, single-cell samples are automatically pooled into sets and transferred into a 384-well plate, which is then placed into the autosampler for automated injection for LC-MS/MS. Any system that supports 384-well plate injection (such as Dionex 3000) can implement this workflow.

**Figure 2.**
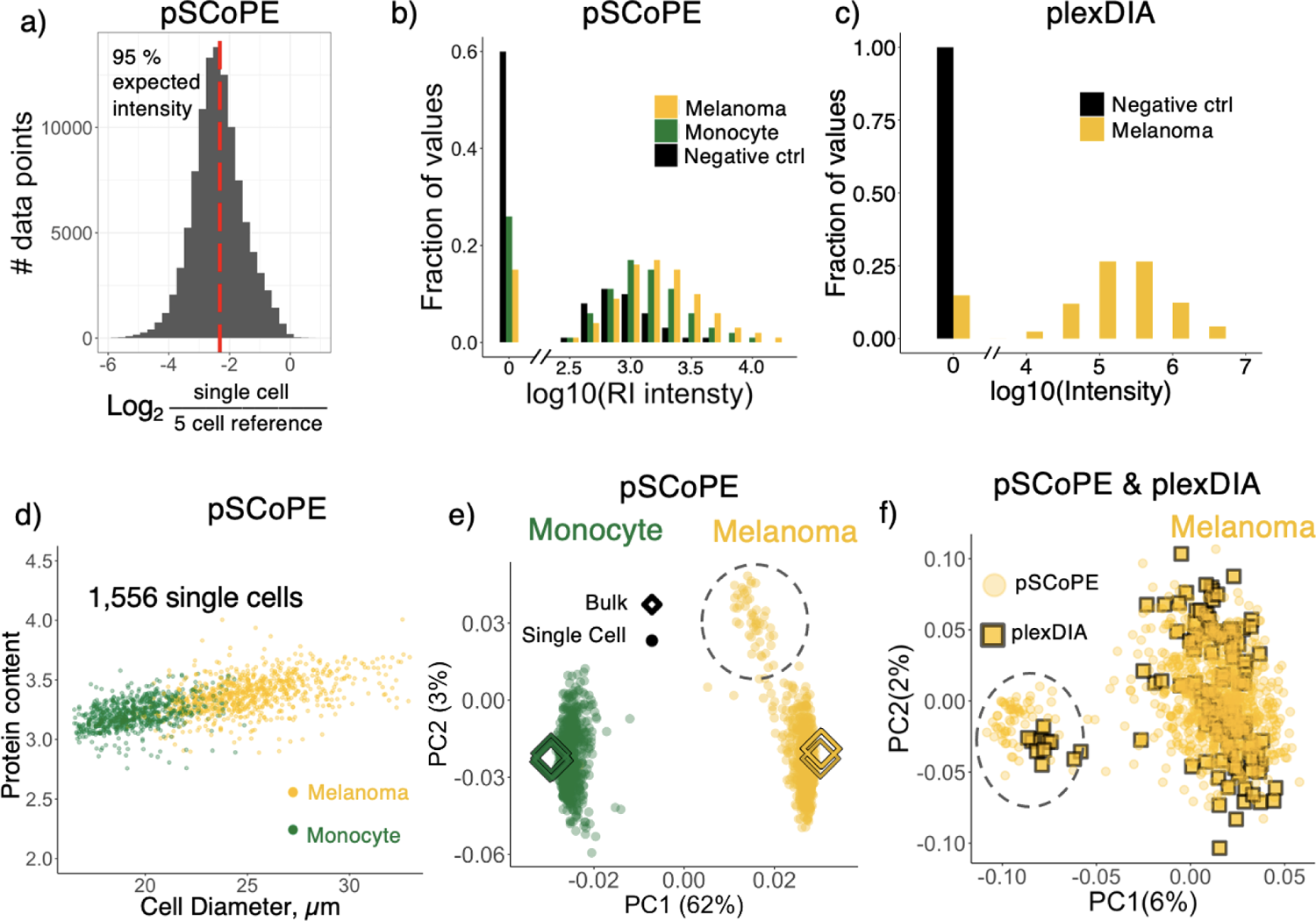
Single-cell data quality control and exploration. (**a**) The efficiency of delivering proteins from single cells is estimated as the distribution of ratios between reporter ions from single cells and 5-cell reference samples. Single cell intensity shows 95 % of the expected 1:5 ratio denoted by the red dotted line. The data were acquired by a Q exactive using pSCoPE with 100-cell isobaric carriers. (**b**) Distributions of reporter ion intensities from single melanoma, monocyte, and negative controls from the pSCoPE sets^27^. (**c**) MS1 intensity of precursors from single melanoma cells and from negative controls from the plexDIA sets^24^. (**d**) The protein content (estimated as the average reporter ion intensity) is plotted against the diameter of cells estimated from images. The correlation suggests consistent delivery of proteins. (**e**) Principal component analysis (PCA) of single cells acquired using pSCoPE with 100-cell isobaric carriers. The single cells cluster by cell type, melanoma and U937 monocytes. Bulk samples of 200 cells labeled with TMTpro are also projected onto the PCA to evaluate the agreement between relative protein quantification in bulk and single-cell samples. A separate smaller cluster of melanoma cells is enclosed in a dotted circle. (**f**) Biological replicates of melanoma cells analyzed by pSCoPE or by plexDIA were jointly projected by PCA. The projections show alignment of two observed sub-clusters; the quantitative agreement is shown in Extended Data Fig. 9b.

To facilitate this flexible design, we needed reagents, compatible both with analysis by LC-MS and our surface design. To this end, we introduced the use of 100 % dimethyl sulfoxide (DMSO) as a reagent for cell lysis and protein extraction. Its low vapor pressure enables nanoliter droplets to persist on the surface of the glass slide. Furthermore, its compatibility with MS analysis obviates sample cleanup and the associated losses and workflow complications. Control experiments indicate that DMSO efficiently delivers proteins to MS analysis without detectable bias for proteins from different cellular compartments (Extended Data Fig. 3a,c) and supports accurate relative protein quantification (Extended Data Fig. 3b,d).

**Figure 3.**
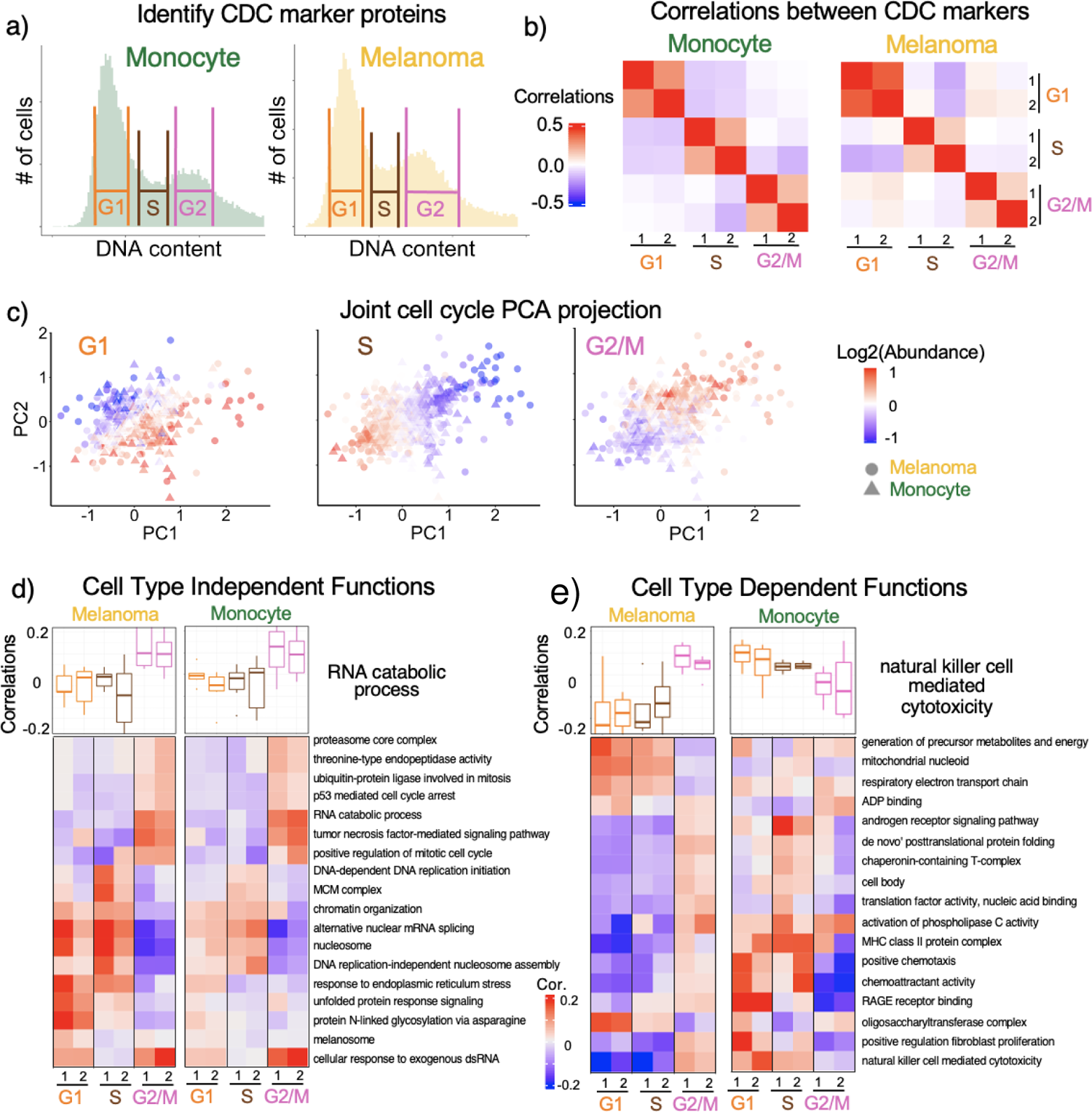
Identifying functional sets of proteins that covary with CDC-markers. (**a**) Distributions of DNA content for FACS-sorted cells used to identify proteins whose abundance varies with CDC phase. (**b**) Correlations between CDC protein markers computed within the single cells from each type in the pSCoPE dataset. (**c**) PCA of melanoma and monocyte cells in the space of CDC periodic genes. Cells in each PCA plot are colored by the mean abundance of proteins annotated to the indicated phase. (**d**) The boxplots display distributions for correlations between the CDC-phase markers and proteins from the RNA catabolism protein set. The correlations were computed for single cells randomly partitioned into two non-overlapping subsets (marked by 1 and 2) to explore their reproducibility. The difference between the distributions for a protein set was evaluated by 1-way ANOVA analysis to estimate statistical significance. The distributions for other protein sets that significantly (FDR *<* 5%) covary in a similar way between the two cell lines are summarized with their medians plotted as a heatmap. (**e**) Similar analysis and display as in panel d was used to visualize protein sets whose covariation with the CDC differs between the two cell types.

Having benchmarked cell lysis and protein extraction by DMSO, we integrated it into the nPOP workflow by printing an experimenter-defined array of DMSO droplets into which single cells are subsequently dispensed for lysis, Fig. 1c. After lysis, proteins are digested for 4 hours by adding 12 *µl* of digestion buffer containing trypsin protease and HEPES. The final concentration of trypsin in the droplet is 90 ng/*µl* along with 4 mM HEPES pH 8.5. The small droplets allow us to reduce the overall amount of trypsin to 1.8 ng (6-fold lower than plate based sample preparation^25^) while greatly increasing concentration of trypsin. This allows for efficient and consistent digestion of peptides, resulting in about 5 *−* 10% missed cleaved rate across the single cells, Extended Data Fig. 5a,b. The reduced amounts of reagents also contributes to reduced contaminant ions, as reflected in the fact that singly charged ions are over 10-fold less abundant than the peptide ions, Extended Data Fig. 4a. In fact, nPOP is the only nanoliter sample preparation that has been utilized without sample cleanup, Extended Data Fig. 2.

**Figure 4.**
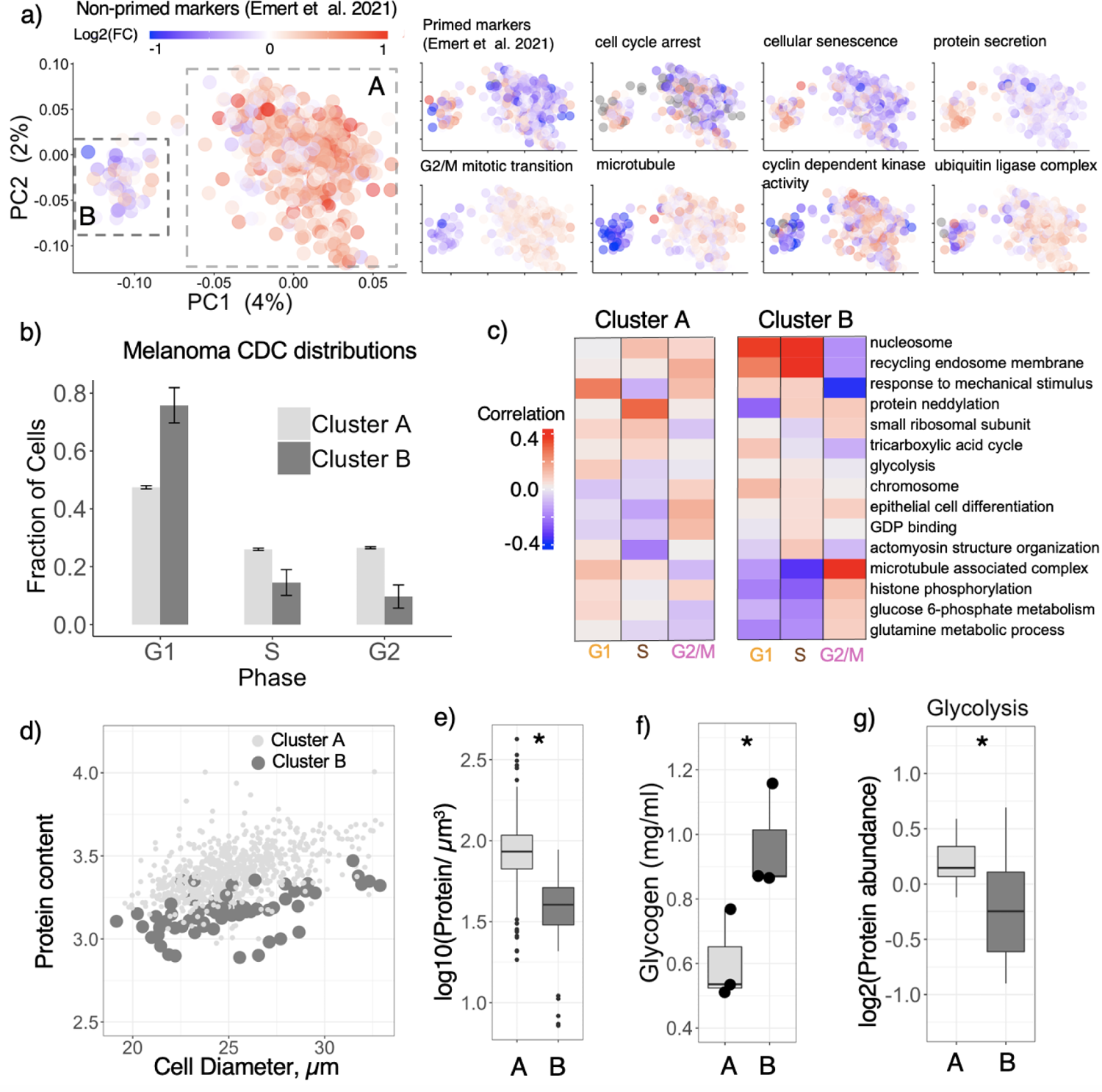
Proteomic, cell-cycle, and metabolic differences between melanoma clusters. (**a**) Single melanoma cells projected by PCA and colored based on the mean abundance of proteins whose transcripts were previously identified as markers of primed cells by Emert *et al.*^7^. The single cells were also colored by the average abundance of protein sets exhibiting significant (*FDR <* 1%) enrichment in cluster A (bottom row) or cluster B (top row). (**b**) Distributions of cells by CDC-phase for cells from cluster A and B. CDC-phases were determined from marker proteins from Fig. 3. (**c**) Protein sets showing distinct covariation in clusters A and B. The analysis and display are as in Fig. 3e. (**d**) Mean reporter ion intensity versus cell diameter for melanoma cells. (**e**) The asterisk (*∗*) indicates *p <* 0.05. Protein concentrations for cells form cluster A and cluster B as defined by total reporter ion signal divided by cell volume. (**f**) Glycogen content measured from 10,000 cells from cluster A or cluster B. (**g**) Glycolytic enzymes have higher abundance in cluster A.

**Figure 5.**
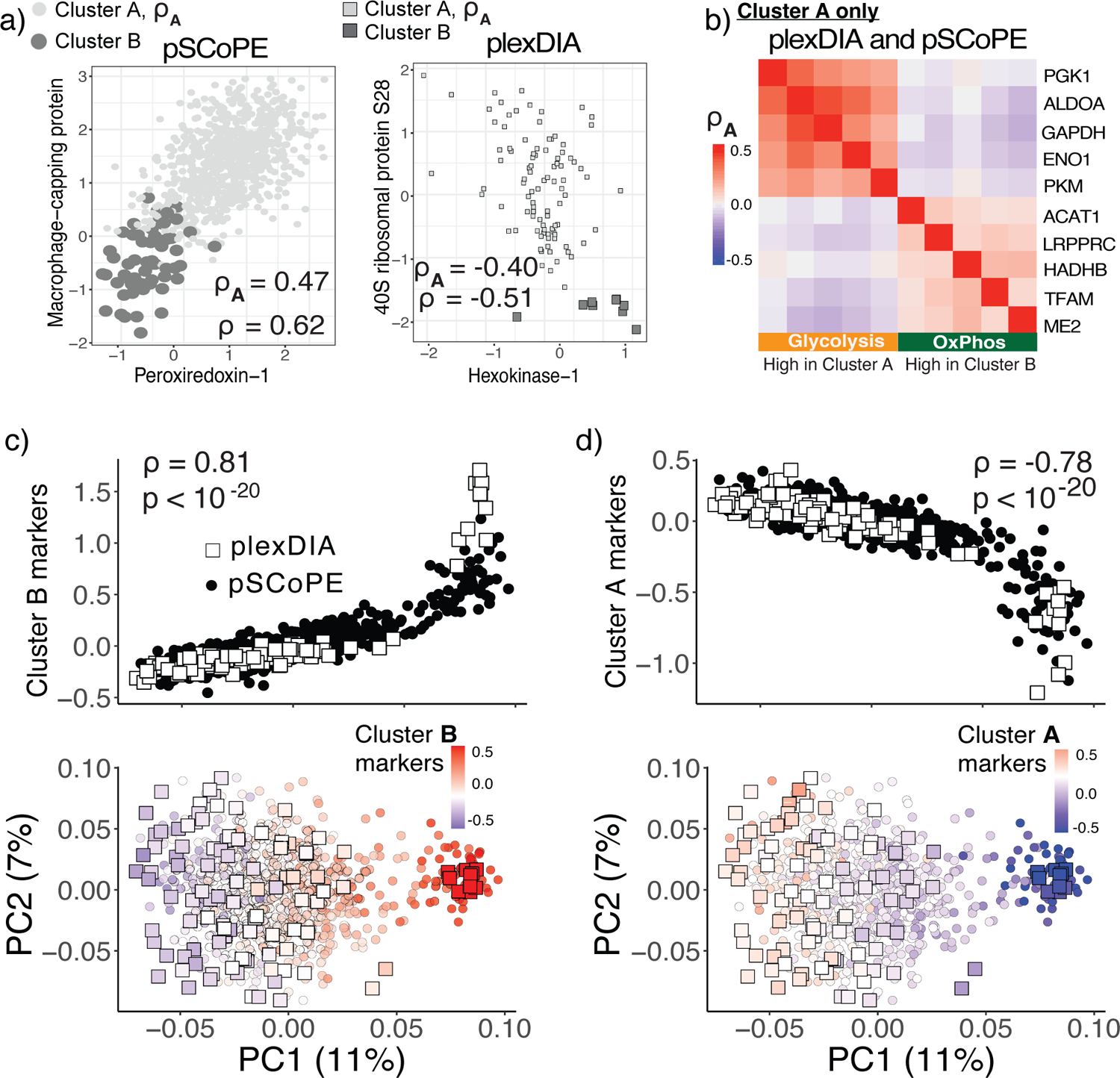
Metabolic enzymes and ribosomal proteins covary and exhibit abundance gradients within the non-primed melanoma sup-population. (a) Scatter plots of protein abundances across single cells for proteins identified to be differentially abundant between cluster A and B in the integrated data set of pSCoPE and plexDIA measurements. (b) Correlations between proteins computed within cluster A. Glycolytic proteins (which are upregulated in cluster A) are positively correlated to each other and inversely correlated to proteins participating in oxidative phosphorylation (which are upregulated in cluster B). Correlations are computed only from cells assigned to cluster A. Exploration of all differentially abundant proteins between cluster A and cluster B reveals a gradient of states for proteins up-regulated in cluster A (c), and for proteins up-regulated in cluster B (d).

To control evaporation throughout digestion, chamber humidity is maintained at 75%, and slide temperature is maintained at 1 *°C* below the dew point. Furthermore, we dispense a perimeter of water droplets around the samples, Fig. 1b. These measures allow droplets to maintain their size and volume throughout as they sit on the slide for several hours through incubation, such as protein digestion, Extended Data Fig. 1.

If choosing to label peptides for multiplexed analysis, we found that the commonly used approach of dissolving labels in acetonitrile was unreliable with our platform’s dispensing method due to the reagent’s low density and surface tension. By substituting DMSO for acetonitrile as the label solvent, we observed robust dispensing for picoliter dispense volumes over hundreds of samples. We validated this approach by measuring labeling efficiency from many single cells pooled together and found that 99 % of all available n-termini and lysine residues were TMT labeled, Extended Data Fig. 5c.

Finally, we use the CellenONE nozzle to sequentially pool clusters of labeled single cells that comprise a set into a single pool, aspirates it, and dispenses it into a 384 well plate in a fully automated fashion, Fig. 1c. A video of this process can be observed in supplemental video 3. The nozzle is washed for 10 seconds in between each sample collection. The 384-well plate can then be placed in a speed vacuum to evaporate any excess liquid, and then sealed with foil for storage at −80 degrees Celsius. Before injecting samples for analysis, wells can be rehydrated with 0.1 % formic acid (which may contain isobaric carrier diluted to the desired concentration), resealed with foil, and placed into a compatible autosampler for automated sample injection and MS analysis, Fig. 1d.

Hands on time for preparing about 2,000 single cells by nPOP is approximately 4 hours, which includes dispensing the initial DMSO droplets, about 2,000 single cells, and trypsin takes about 3 hours. Dispensing for labeling and quenching takes about 1 hour. Hands off time for digestion is 4 hours and sample pickup is about 45-55 seconds per sample. To obviate a long day of sample preparation, a pause point can be added after digestion: digested peptides are dried out on the slide and left to sit in the enclosed environment overnight. The next day samples can be re-hydrated for labeling, quenching, and subsequent collection (see methods).

Taking advantage of the flexibility afforded by nPOP, we quantified proteins with two different methods, one performing quantification at MS1 level and the other at MS2 level. Specifically, using nPOP we prepared multiplexed single-cell samples for data independent acquisition by plex-DIA^24, 26^ and for prioritized data acquisition by pSCoPE^27^, Fig. 2. These methods have different biases and thus can support cross-validation of protein quantification. The pSCoPE samples were prepared according to the SCoPE2 design with TMTpro 18-plex labels^14, 25^. The isobaric carrier and reference samples corresponded to 100-cell (supplemental figure 8, S1) and 5-cell sample loads. For pSCoPE, we prepared 2,016 single-cell samples (using layout AL-01 and including both nehative controls and single cells) in one day and successfully analyzed 1,556 single cells, Fig. 2c. This is about 6-fold higher than previously recorded for a single sample preparation, and over 7 fold higher than other small volume methods, Extended Data Fig. 2. This dataset will herein be referred to as pSCoPE. MS analysis used 0.5 Th isolation windows for MS2 scans, which resulted in high spectral purity, as seen from a DO-MS report highlighting this and other aspects of the LC-MS analysis (supplemental data 5). This is lower than the 2,016 capacity due to: 1) including 128 negative controls (droplets that did not receive single cells), 2) having 175 single cells excluded from analysis and 3) 15 sets lost because of LC malfunctions. Each set contains 6-7 monocyte and 6-7 melanoma cells that are randomly distributed amongst the different TMT channels, see Metadata (supplemental data 4). Using the plexDIA design, we prepare a biological replicate of an additional 768 melanoma cells in one day, and analyzed 44 sets having 104 single cells and 9 negative controls. This dataset will herein be referred to as plexDIA. Processed protein by single cell matrices for pSCoPE and plexDIA can be found in supplemental data file 9. All samples were analyzed on a Q-Exactive Basic mass spectrometer.

Major objectives for nPOP included efficient delivery of proteins to MS detectors and low background noise. These factors directly evaluate sample preparation, and thus we sought to rigorously quantify them, Fig. 2a-c. To evaluate efficiency of protein delivery, we compared reporter ion intensities in single cells to the corresponding reporter ion intensities of the 5-cell reference, which was prepared in bulk. The results indicate that single cell intensities are on average 95 % the expected 1:5 ratio, Fig. 2a, suggesting that nPOP delivers proteins from single cells at 95 % of the efficiency of optimized bulk sample preparation, which suffers less from adhesion related losses.

To quantify the level of background noise, we evaluated the raw reporter ion intensities in negative controls, Fig. 2b,c. Negative control droplets are treated identically as single-cell droplets with the sole exception that no single cells are added. Thus, MS intensity measured in these samples reflects background contamination that may arise from sources such as reagent impurities, cross labeling between single cells, or interference between ions. The background intensities in negative controls of pSCoPE sets are not detected for 60% of the peptides and mean intensity per negative control is about 20-fold lower than the mean single cell intensity, Fig. 2b. This signal is entirely absent in the plexDIA measurements, Fig. 2c. These results suggest that labeling is specific and efficient for singe-cell samples prepared with nPOP, though the co-isolation of labeled contaminant ions can contribute to background signal when using isobaric mass tags.

A less direct measure of sample preparation is the number of confidently identified peptides, as these numbers depend on the MS instrument used, length of gradient, and data acquisition method and type, cell size, and ion accumulation times. Nonetheless, the proteins quantified per single cells of our samples prepared by nPOP exceeds significantly our previous results based on isobaric carrier^14, 25^, though acquisition methods differed as well. Specifically, the pSCoPE data set quantified on average 997 proteins per cell, with 2,844 proteins quantified across the 1,556 single cells prepared by nPOP, Extended Data Fig. 6a. In the plexDIA biological replicate data set, we quantified an average of 410 proteins per cell, Extended Data Fig. 6d.

The distributions of intensities for single cells show that peptides from melanoma cells are more abundant than peptides from monocyte cells, possibly reflecting the different cell sizes Fig. 2b. To further test the extent to which higher reporter ion signal in the melanoma cells reflects larger cell size, we plotted the measured diameter for single cells against the average reporter ion signal, Fig. 2d. The correlation between cell diameter and average intensity, *ρ* = 0.81, indicates that nPOP consistently delivers proteins to MS analysis without significant variations from cell to cell. This correlation between protein abundance and cell size places an upper bound on the variability of protein delivery by nPOP. Deviations from perfect correlation may also reflect differences in protein concentrations in individual single cells (as demonstrated below) or errors in estimating cell size, either because of errors in measuring cell diameters or deviation of cellular shapes from spherical shapes.

As an additional QC metric, we evaluated the agreement in variability in protein quantification derived from different peptides originating from the same protein. This variability is quantified by the coefficients of variation (CVs) and the average protein CV is significantly lower for the single cells than the negative controls, Extended Data Fig. 6a,c. Furthermore, the small spread of the CV distributions suggests high consistency of the automated sample preparation technique, and an improvement over the variability observed with previous sample preparation methods^25^.

Next, we performed principal component analysis (PCA) of the single-cell protein dataset using all 2,844 quantified proteins by pSCoPE, Fig. 2e. To compute the PCA, missing values were imputed using k-nearest neighbors imputation. The PCA of single cells analyzed by pSCoPE identified three distinct clusters of cells, one corresponding the monocytes, and two sub-populations of melanoma cells. The first principal component (PC1) separates monocytes and melanoma cells and accounts for 62 % of the total variance in the dataset. To evaluate whether this separation reflects technical artifacts, such as differences in cell size or missing data, or biological differences between cell types, we projected the proteomes of 200-cell samples of melanoma and monocyte cells analyzed by established bulk methods, Fig. 2e. The results indicate that the clustering of single cells is fully consistent with the protein levels measured from the bulk samples and not correlated to artifacts, such as the slide number, variability in digestion or the position on the slide, Extended Data Fig. 7. As an additional check that separation was not caused by differences in culture conditions shared between bulk and single cells, we colored single cells by the average abundance of proteins belonging to protein sets related to melanoma or monocyte biology, Extended Data Fig. 8a,b. Several protein sets associated with melanoma such as epidermis development, epithelial cell proliferation, and melanosome were associated with melanoma cells (all *p <* 10*^−^*^15^). Protein sets including defense response to bacterium, MHC class I, and leukocyte chemotaxis were associated with monocyte cells (all *p <* 10*^−^*^15^).

In addition to quantifying differences between monocyte and melanoma cells, we also observed two distinct populations within the melanoma cells, Fig. 2e. To validate this observation, we prepared an additional biological replicate, analyzed by an orthogonal mass spectrometry quantitation method, plexDIA, which performs precursor quantitation at the MS1 level^24^. When projecting melanoma cells analyzed by plexDIA into a low dimensional space via PCA, we similarly noticed two distinct clusters separated on PC1, Extended Data Fig. 9a. We next sought to see if these two clusters corresponded to the same cellular populations by comparing protein fold changes between the two clusters, Extended Data Fig. 9b. The strong agreement observed between these different measurements, *ρ* = 0.70, suggests that the subpopulations in Fig. 2e are defined by protein quantification that is reproducible across biological replicates and very different MS methodologies. Thus, we projected these two data sets into the same space via PCA, Fig. 2f, and the alignment of the clusters further confirmed the concordance between the melanoma populations.

### Protein covariation with the cell cycle

Next, we moved on to the more challenging problem of quantifying CDC-related protein covariation within a cell type in the pSCoPE data set containing monocyte and melanoma cells. As a first step, we evaluated the potential to classify individual cells by their cell cycle phase. To obtain a list of proteins whose abundance varies periodically with the cell division cycle, we first sorted populations of each cell type based off their DNA content, Fig. 3a. We quantified the proteomes of the sorted cells and identified proteins whose abundance differs in G1, S, and G2/M phase for both cell types. We identified a set of 596 proteins with significant (FDR = 1%, supplementary table 6) changes in abundance across the CDC phases, including many canonical CDC markers, Extended Data Fig. 10a.

Using this set of CDC periodic proteins, we sought to construct CDC markers that robustly represent cell cycle phases in our pSCoPE dataset. To this end, we used a systematic cross-validation procedure to select the subset of proteins showing CDC-periodicity in the bulk data (the proteins with the highest fold-change between each CDC phases) and exhibiting the expected correlation pattern within the single-cell data: namely, markers from the same phase should correlate positively to each other while markers from different phases should correlate negatively to each other, Fig. 3b. Each CDC marker vector represented the average profile of several proteins whose abundances peak in the same CDC phase. These average profiles were chosen to average out measurement noise for individual proteins and to increase robustness.

For additional validation, we constructed the CDC markers using only the monocyte cells and tested them on the melanoma cells, Fig. 3b. Specifically, we evaluated whether CDC markers selected from the monocyte dataset exhibit the expected correlation pattern in the melanoma dataset that was not used for the marker construction, Fig. 3b. The correlation (*ρ* = 0.81*, p* = 0.00052) between the two patterns cross-validates the chosen markers. Having validated the protein markers, we averaged protein markers for the same phase, creating one marker for each CDC phase to be used in downstream analysis. The list of proteins comprised of each marker can be found in supplemental table 6.

To further explore our ability to capture CDC-related protein dynamics, we used principal component analysis to project the proteomes of both melanoma and monocyte cells into a joint 2-dimensional space of the CDC marker proteins. Each cell was then color-coded based on the mean abundance of a protein marker for the indicated CDC phase, Fig. 3c. The cells from both cell types cluster by CDC phase, which further suggests that the data capture CDC-related protein dynamics. Lack of discreetness of the cells on the edges between different regions of the PCA may reflect that the cell cycle is a continuous process and these cells may be between two different phases.

Next we focused on identifying proteins that covary with the CDC periodic markers. To identify such covariation, the phase marker vectors were correlated to the measured protein abundances (only observed values) of all proteins with at least 150 observations quantified across the single cells. Hundreds of proteins (381 for melanoma and 292 for monocyte cells, supplemental table 6) are significantly (FDR *<* 5%) correlated to the CDC markers, suggesting that these proteins are CDC periodic (supplemental table 6). The list includes well characterized CDC proteins such as Mitotic checkpoint serine/threonine-protein kinase BUB1, which facilitates spindle-assembly. BUB1 is positively correlated with the G2 markers in both melanoma and monocyte populations, *p <* 10*^−^*^5^, *p <* 10*^−^*^4^, respectively.

To increase the statistical power and identify functional covariation with the CDC, we next focused on the covariation of phase markers and proteins with similar functions as defined by the gene ontology (GO). We restricted this analysis to proteins with at least 150 observations, resulting in set of 1,551 proteins with about 60% values present in the cell x protein matrix. For each protein set, we computed the correlations between its proteins and each CDC-phase marker vector. The distributions of these correlations (as shown with the boxplots for RNA catabolism in Fig. 3d) were then compared using ANOVA to evaluate the statistical significance of differences between cell cycle phases. To evaluate the reproducibility and reliability of these correlations, we randomly partitioned the single cells into two non-overlapping subsets of cells and displayed the correlations estimated for each subset, marked by 1 and 2 in Fig. 3d. The correlations estimated from these non-overlapping subsets are similar (correlation *ρ >* 0.8), which indicates that the analysis captures systematic CDC-related protein covariation. For RNA catabolism, the distributions of correlations differ significantly between the CDC phases, and this phase-specific covariation is similar for the two cell types. Similar to RNA catabolism, other protein sets show covariation with the phase markers that is similar for the two cell types, and instead of displaying the boxplot distributions for all of them, we summarized the distributions of correlations with their means displayed as a heatmap, Fig. 3d. Such functions with shared covariation include proteolysis in G2/M phase, which is consistent with the role of protein degradation in cell cycle progression^28^. Additionally, terms related to DNA repair and translation are correlated with G1 markers suggesting the role of cell growth and DNA repair post mitosis. The majority of the 117 significant protein sets (supplemental table 6) showed concerted trends between the two cell types, highlighting the conservation CDC related processes. In addition to finding groups of proteins that show similar cell cycle covariation between cell types, several protein sets co-varied differentially between the cell types, Fig. 3e. They include terms related to cell signalling, metabolic, and immunological processes, which may reflect the role of the monocyte as an immune cell.

The correlation analysis may detect variation in protein abundance not between phases, but also within a phase. This higher time resolution cannot be validated by our data from cells sorted based on DNA content (Extended Data Fig. 10a,b). However, the correlation differences between CDC phases (Fig. 3e,d) can be cross-validated with the bulk DIA data. To perform such validation, we compared the protein sets with significant marker correlations to the protein sets with significant fold-changes measured in the cells isolated based on DNA content. The results (Extended Data Fig. 10b) indicate significant (*p <* 0.05) similarity between the protein dynamics estimated from single-cell covariation and from the classical protein abundance measurements, and thus further validate our protein covariation analysis.

### Protein covariation within melanoma sub-populations

We next turned our attention to the two distinct clusters of melanoma cells observed in the pSCoPE data, Fig. 2e,f. Recent studies of these melanoma cells identified a transcriptomic signature associated with a cell state that is more likely to resist treatment by the cancer drug vemurafenib^7, 29, 30^. We estimated the differential protein abundance of these markers, which is visualized by color coding the cells by the mean abundance of proteins whose transcripts were identified as markers for each cell state^7^. The results indicate that the larger cluster (cluster A) have higher abundance of the non-primed markers (*p <* 10*^−^*^1^^5^) while the smaller cluster (cluster B) have higher abundance of the primed markers (*p <* 10*^−^*^5^), Fig. 4a. These results establish a link between the clusters in our data and priming for drug resistance.

Having established correspondence between the populations, we sought to identify additional protein differences between the two clusters by performing Protein Set Enrichment Analysis (PSEA). It resulted in 84 sets of functionally related proteins exhibiting differential abundance between the clusters at FDR *<* 1% and an effect size *>* 50%, (supplemental table 7). A selection of these results are displayed in Fig. 4a, in which cells are color-coded by the median abundance of the proteins associated with each protein set. Protein sets related to G2/M transition of mitosis, cyclin-dependant kinase activity and protein degradation are more abundant in cluster A. In contrast, protein sets with increased abundance in cluster B are related to senescence and cell-cycle arrest. These results suggest that cells in cluster A are more proliferative than those in cluster B, consistent with a prior report^29^.

To further explore CDC differences between clusters A and B, we quantified the distribution of cells in each CDC phase across the two clusters. The results in Fig. 4b reveal a substantial difference: The majority (78 %) of cells in cluster B are in G1 phase while only 4 % of its cells are in G2 phase. In contrast, cells of cluster A are more evenly distributed across all phases, Fig. 4b. This result further bolsters the conclusion that cells in cluster B divide slower than those in cluster A. We next sought to identify functional groups of proteins that may co-vary differently with the cell cycle in the two clusters. Upon repeating the analysis from Fig. 3e on the two melanoma clusters, we found several sets of proteins that correlated significantly to the CDC markers but differed between clusters A and B, Fig. 4c. Specifically, positive regulation of cell growth is strongly correlated to G1 markers in population A but not in cluster B, consistent with slower proliferation of population B. These differences in protein dynamics between clusters A and B suggest specific processes that might be regulated differently between the clusters and potentially contribute to the drug-resistance priming. These processes shown in Fig. 4c include carbohydrate metabolism (glycolusis, TCA cycle, glucose 6-phosphate metabolism) and protein synthesis (ribosomes).

### Glycogen accumulation in primed melanoma cells

We next sought to examine any additional phenotypic features of the identified sub-population. One readout available from nPOP includes images of the single cells which allow estimating their cell diameters. When comparing the relationships between cell diameters and estimated total protein content, we noticed a significant offset between clusters A and B, Fig. 4d. This offset might reflect incomplete extraction of protein from cluster B, though this possibility seems unlikely given the efficient lysis and protein extraction observed upon addition of DMSO, Extended Data Fig. 3. Alternatively, it may be due to a significant (*p <* 10*^−^*^15^) difference in total protein concentration per unit cellular volume, Fig. 4e.

If protein concentration is lower in cluster B cells, another cellular component must have higher concentration. We hypothesized that this component is likely glycogen since accumulation of glycogen has been previously observed in G1 phase and in slowly dividing cells^31, 32^. To test this hypothesis, we sorted 30,000 cells from each of the two clusters by FACS based on the abundance of CD49c, a priming marker reported by Emert *et al.*^7^ and associated with cluster B in our single-cell proteomics data. Specifically, cells from cluster B were isolated as the population with top 1 % expression of CD49c, and cells from cluster A were isolated as the population with 25% to 75% expression range of CD49c, see supplemental figure 8, S4. After sorting equally sized populations, we performed an enzymatic assay to quantify total glycogen amount in each population. The results in Fig. 4f showed a significant (*p <* 0.05) increase of *∼* 50% in glycogen for cluster B cells. This is consistent with our hypothesis that increased glycogen storage is responsible for the reduction in protein concentration in cluster B cells. To further evaluate and extend the difference in glycogen metabolism, we test for functional enrichment of glucose metabolic pathways. This analysis showed that proteins involved in glycolosis, the catabolism of glucose for energy, were upregulated in cluster A, which is consistent with the reduced amount of measured glycogen, Fig. 4g.

### Gradients within the non-primed melanoma cluster

Next, we sought to evaluate the degree of coordinated expression of proteins associated with either cluster B or cluster A. Co-expression of transcripts associated with drug priming has been previously reported in “jackpot cells”, which results in much higher frequency of these cells compared to the expected frequency of expression was not correlated^33^. Consistently, we observed that many genes are co-expressed (covary) at the protein level *between* clusters A and B, Fig. 4. Yet, it remains unclear whether proteins covary *within* a cluster. We evaluated this possibility for the 75 proteins that significantly (FDR *<* 1%) change in abundance between clusters A and B by 2 fold or more in the combined dataset shown in Fig. 2f; The proteins are listed in supplemental table 7. Pairs of these proteins were directly examined for covariation both within and between clusters, Fig. 5a. The results indicate that protein pairs such as 40s ribosomal protein S28 and Hexokinase-1 covary significantly not only across the clusters (*ρ* = *−*0.51, *p <* 10*^−^*^15^), but also within cluster A, (*ρ* = 0.40, *p <* 10*^−^*^7^).

To extend this analysis beyond protein pairs, we examined functional groups of proteins correlated within the cluster A of non-primed melanoma cells. To avoid detecting covariation that is due to systematic bias of a measurement method, we computed the correlations using the combined pSCoPE and plexDIA dataset, Fig. 5b. The results indicate positive correlations within the clusters of glycolitic enzymes and within the cluster of oxidative phosphorylation proteins and negative correlation between the clusters. This pattern suggests a gradient in the extent to which non-primed melanoma cells derive their energy aerobically. This within cluster gradient parallels the the differential abundance of these protein sets across clusters A and B (Fig. 5b).

To systematically explore this parallel for all 75 proteins identified as differentially abundant (FDR *<* 1% and an effect size *>* 2 fold) between clusters A and B, we performed PCA in the space of these proteins, Fig. 5c,d. The results indicated not only the expected separation between clusters A and B but also a gradient of changes in protein abundance within cluster A, Fig. 5c,d. This gradient generalizes the observation from the protein covariation pattern in cluster A (Fig. 5b) and extends is to the entire set of proteins that are differentially abundant between cluster A and B. The agreement between pSCoPE and plexDIA measurements suggests that this gradient does not reflect method specific biases but rather reflects concerted biological variability. It also suggests that some cells in cluster A are more likely to transition to the state associated with priming represented by cluster B.

## Discussion

Existing single-cell omics methods excel at classifying cells by cell type. However, the regulatory dynamics resulting in cell-to-cell variability within a cell type are more challenging to analyze^5^. To support such analysis, we introduce a highly parallel sample preparation method, nPOP, that allowed us to simultaneously prepare over 1,500 single cells. This demonstrated throughput can be further increased by increasing the density of droplets and the number of slides (Fig. 1), which will support the projected increase in the throughput of single-cell proteomics^10, 26^. nPOP reduced background noise (Fig. 2 and Extended Data Fig. 4) and increased consistency of single-cell proteomic sample preparation, Extended Data Fig. 5 and Extended Data Fig. 7. The resulting increases in throughput and data quality enable high-throughput, high-power biological analysis demonstrated here with the analysis of cell cycle and melanoma drug resistance. These benefits will extend to many other systems^10, 34^. While here we demonstrate nPOP using CellenONE, we expect that nPOP can be implemented on additional liquid handling platforms.

Important biological observations, such as the gradient of protein abundance in non-primed melamona cells, were demonstrated by two very different MS methodologies. This demonstration was crucially enabled by the flexibility of nPOP sample preparation tow work with both 18-plex isobaric mass tags^27, 35^ and 3-plex non-isobaric mass and plexDIA^24, 26^. This flexibility stems from the ease of computationally programming different droplet layouts and workflows, Fig. 1. Thus, it can be easily extended to workflows without labels (label-free analysis) as well as multiplexed analysis with different number of labels, which likely to be very useful for new high-plex MS methods under development^26^. To facilitate all workflows, we developed an detailed protocol^36^.

The throughput and consistency afforded by nPOP allowed us to perform adequately-powered covariation analysis, which identified new proteins and functional groups of proteins associated with the cell cycle without the artifacts of synchronized cell cultures^37^. Unbiased clustering of both plexDIA and pSCoPE data identified (and cross-validated) a sub-population of melanoma cells (Fig. 2) expressing markers of priming for drug resistance (Fig. 4).

Many of our findings depended crucially on performing single-cell measurements and could not have been made if we had isolated and analyzed cell populations from cluster A and B. For example, the gradients and concerted pattern of variation *within* the non-primed cells could not be detected by analyzing the cells from this cluster as an isolated subpopulation. This gradient of states may indicate a path of variation along which cells switch from the non-primed cluster to the cluster associated with priming for drug resistance. These results demonstrate the feasibility of inferring co-regulation of biological processes from single-cell proteomics measurements.

## Data availability

A description of all available data can be found in supplemental table 1. This includes MassIVE submission, supplementary files, and additional processed data provided using community recommended best practices^38^. All raw data and search engine outputs can be found on MassIVE with ID: MSV000089159. Additional processed data, protocols, video tutorials, code for reproducing the analysis and other resources are available at: scp.slavovlab.net/nPOP and scp.slavovlab.net/Leduc et al 2022

**Table 1.** All reagents used for sample preparation with nPOP. Listed are volumes added, concentrations in the dispensing solution, and concentrations in the droplets. ^1^Aqueous TEAB solution is added only for mTRAQ labeling (not for TMT) since it is required for optimal mTRAQ labeling.

|  | Reagent | Dispense conc. | Volume added | Concentration in droplet |
| --- | --- | --- | --- | --- |
| 1 | DMSO | 100 % | 8 nl | 100 % |
| 2 | Cell | 300 cell/μl in 1x PBS | 300 pl | 1 cell |
| 3 | Digest mix |  | 13 nl | 60 % (aq) |
| | Trypsin Gold (aq) | 150 ng/μl | | 90 ng/μl |
|  | HEPES (aq) | 6 mM |  | 4 mM |
| 4 | TMT (DMSO) | 8.3 μg/μl | 30 nl | 8.3 μg/μl |
| 5 | mTRAQ (DMSO) | 8.3 μg/μl | 30 nl | 8.3 μg/μl |
| 6 | TEAB (aq)<sup>1</sup> | 200 mM | 20 nl | 50 mM |
| 7 | Hydroxylamine (HA) | 5 % | 2 x 20 nl | 3 % |

**Table 2.** Specific terms used in describing nPOP sample preparation procedure and their corresponding definitions

| CellenONE terms | Description |
| --- | --- |
| PDC | Piezoelectric dispensing capillary for dispensing cells and reagents |
| Field file | Text file describing position of droplets on glass slide |
| Spot_Run | CellenONE method dispenses current content of PDC to glass slide locations specified by the field file |
| Pickup_Sample_and_Dispose | CellenONE method that automatically picks reagents from 384 well plate and dispenses to specified position |
| CellenONE_Basic | CellenONE method used for isolating and dispensing single cells to specified position |
| Pickup_off_slide | CellenONE method used for collecting samples dispensing into 384 well plate |
| DipWash | Dips nozzle into water to wash off outside |

## Code availability

The code used for data analysis and figures is at: github.com/Slavovlab/nPOP

## Protocol availability

A video step-by-step protocol is available on YoutTube and a detailed step-by-step protocol is available at protocols.io with doi: 10.17504/protocols.io.b67erhje.

## Acknowledgments

We thank A. Chen for early experiments with the organic solvent lysis, A. Murphy for assistance with using CellenONE, and H. Specht for discussions and constructive comments. This work was funded by a New Innovator Award from the NIGMS from the National Institutes of Health to N.S. under Award Number DP2GM123497, an Allen Distinguished Investigator award through the Paul G. Allen Frontiers Group to N.S., a Seed Networks Award from CZI CZF2019-002424 to N.S., through a Merck Exploratory Science Center Fellowship, Merck Sharpe & Dohme Corp. to N.S. Funding bodies had no role in data collection, analysis, and interpretation.

## Correspondence

Correspondence and requests should be addressed to

## Author Contributions

Experimental design: A.L., G.H. and N.S. Single-cell LC-MS/MS: A.L., G.H. Sample preparation: A.L, S.K, J.C Raising funding & supervision: N.S. Data analysis: A.L., S.K. and N.S. Writing & editing: A.L. and N.S.

## Methods

### Sample preparation

#### Cell culture

Jurkat cells were grown as suspension cultures in RPMI medium (HyClone 16777-145) supplemented with 10% fetal bovine serum and 1% penicillin-streptomycin. U-937 cells were grown as suspension cultures in either standard RPMI medium, or heavy SILAC RPMI medium supplemented with +10*Da* Arg and +8*Da* Lys. Cells were passaged when a density of 10*e*6 cells/ml was reached, approximately every two days.

The melanoma cells (WM989-A6-G3, a kind gift from Sydney Shaffer, University of Pennsylvania) were grown as adherent cultures in TU2% media which is composed of 80% MCDB 153 (Sigma-Aldrich M7403), 10% Leibovitz L-15 (ThermoFisher 11415064), 2 % fetal bovine serum, 0.5% penicillin-streptomycin and 1.68mM Calcium Chloride (Sigma-Aldrich 499609). Cells were passaged at 80% confluence (approximately every 3-4 days) in T75 flasks (Millipore-Sigma, Z707546) using 0.25% Trypsin-EDTA (ThermoFisher 25200072) and re-plated at 30% confluence.

#### Preparation of lysis validation samples

To test the extraction efficiency of DMSO lysis, U-937 cells at a concentration of 10,000 cells/*µl* were lysed by incubation with either 6M Urea or 90 % DMSO for 30 minutes each. Two replicates were prepared, one with light SILAC cells lysed with Urea and heavy SILAC cells lysed with DMSO and a label swap of these SILAC conditions. Solutions were then diluted down to a concentration of 1,000 cells/*µl* to dilute urea and DMSO concentrations to 0.6M and 10%, respectively. Suspensions were then combined for digestion. TEAB at pH 8.5 was added to a concentration of 100 mM and trypsin was added to a concentration of 10 ng/*µl*. The mixture was digested overnight at 37 degrees C.

We next sought to measure if systematic artifacts that arise when performing relative quantitation between samples lysed with DMSO. To this end, we compared protein fold changes between Jurkat and monocytes lysed with either DMSO or urea. Heavy SILAC monocytes and light SILAC Jurkat cells were washed and re-suspended in PBS at 10,000 cells per *µl*. Two solutions of 10 *µl*, each containing equal cell count of Jurkat and U-937 cells were made mixed in 1:1 ratios. Samples were lysed in either 90 % DMSO or urea for 30 minutes, diluted 10 fold in HPLC water, and digested as described above. Each sample was then desalted using C18 stage tips.

### Bulk melanoma and monocyte samples for carrier and single cell validation

Cell pellets of 100,000 monocyte (light SILAC, i.e. normal) and melanoma cells were suspended in 50 *µl* of mass spectrometry grade water for a concentration of 2000 cell per *µl*. Samples were lysed and digested via mPOP sample preparation^18^. Briefly, samples were frozen to −80 and then heated to 90 C for 10 minutes for lysis. Trypsin, benzonase, and TEAB buffer were added to concentrations of 10 *µl*, 1X, and 100 mM for overnight digestion at 37C, respectively. Digested samples were then labeled with TMTpro 18-plex labels.

For the bulk validation of single cell data sample, three TMT channels were used as replicates for each cell type. Each TMT channel contained 20 ng of digested amounts of either monocyte or melanoma cells for a final injection amount of 120 ng.

For preparing our isobaric carrier, aliqots consisting of a 1:1 mixture of melanoma and monocyte digested material were labeled with either 126C or 127N. The final diluted concentration was 50 cells of each cell type per *µl* of 126C label (carrier) and 2.5 cells of each type per *µl* of 127N (reference). 1 *µl* of the carrier/reference suspension was injected with each single cell set for analysis.

#### Single cell sample preparation and experimental design

All data files required to prepare samples for nPOP and a direct step by step protocol are available in supplementary files 1 and 11 respectively. Additional resources including a video tutorial for utilizing the CellenONE software for nPOP can be found at scp.slavovlab.net/nPOP.

nPOP reactions are carried out on the surface of a fluorocarbon coated glass slide. Positioning of droplets in the slide is thus adjustable and only limited by the 5 micron increments in which the robot arm can move. In this paper, we demonstrate two potential droplet layouts. These layouts serve different multiplexing schemes, isobaric labeling with TMTpro 18-plex utilizing an isobaric carrier (SCoPE2), and isotopologous mTRAQ for 3-plex plexDIA single cell analysis without the use of carrier. Droplets can be placed anywhere over a regular grid with spaces between droplets separated by 200 microns. In the nPOP protocol, droplets where single cells are prepared are arranged in clusters, with the number of droplets per cluster equals the number of single cells per labeled set. We demonstrate two different droplet design schemes.

**Scheme 1** was designed for TMTpro 18-plex. In this design, a carrier and reference sample are used and two TMT tags, adjacent in mass of reporter ion to the carrier and reference, are left unused because of isotopic carryover from the carrier. This leaves 14 tags for single cells, and thus 14 droplets per cluster. As we analyzed both monocyte and melanoma cells, each set received 6 or 7 of each cell type. With 14 drops per cluster, we fit 36 clusters per slide on 4 different glass slides for 144 sets total.

**Scheme 2** was designed for mTRAQ multiplexed reagents with 3 labels. Here samples are analyzed via DIA and there are 3 single cells per set. No carrier is utilized for these sets. With scheme 2 were able to prepare 64 clusters per slide for 256 overall sets.

For both scheme 1 and scheme 2, droplets within cluster are approximately 700-800 microns apart, and the edge of one cluster to the edge of an adjacent cluster is separated by a distance of 1 mm. Further reducing the space between clusters and droplets can further increase the number of samples per slide for either scheme 1 or scheme 2. We have made several other designs available in supplementary file 1. These include label free and a 10 droplet per set scheme for TMT analysis on timsTOF instruments.

All steps rely on a series of layout files that are appropriated in folders grouped by general choice of droplet layout scheme. For each layout scheme, there are several files for dispensing different reagents. Each time a new reagent is to be dispensed, the corresponding droplet layout file is loaded, which specifies the spaces where each reagent is added on the slide along with the volumes to be added.

Below is a table consisting of all reagents utilized, their stock concentrations, and their concentrations in droplet.

### Detailed steps of nPOP

Below are detailed descriptions for each step of nPOP. Step-by-step protocols are also available on YoutTube and at protocols.io with doi: 10.17504/protocols.io.b67erhje.

1. Setup/usage: Start up CellenONE as instructed by manufacturer. We recommend operating the CellenONE with two PDCs as subsequently described. However the sample preparation is possible with a single standard, medium sized glass PDC. This may come with trade offs in reliability of dispensing. When operating with two PDCs, the first standard, size medium glass PDC is dedicated to handle cell suspensions only. The second size medium PDC with type 2 coating is dedicated for all other reagent handling including organic solvents and protein solutions. After each sample preparation, we recommend that PDCs are washed with ethanol and cleaned under sonication to remove any built up material from inside of the PDC and ensure optimal performance. (15 minutes) Reagents for the sample preparation are loaded into a 384 well plate in volumes of at least 5 *µl* excess for subsequent aspiration and dispensing by the PDCs. Prepare at least 60 *µl* of cells at 300 cell/*µl* in 1x PBS, 500 *µl* of 100 % DMSO, and 60 *µl* of digest mix (150 ng/*µl* Trypsin Gold, 6 mM HEPES) to start the sample prep. (10 minutes)
2. Lysis: Once an array layout scheme is selected, load the field file for the lysis named DMSO L. Load 30 nl of 100 % DMSO into wells D1-D8 of the 384 well plate. Run Pickup Sample and Dispense to spot 8nl of DMSO to each location of the array forming the initial reaction volume for each single cell reaction. Lysis begins when cells are later dispensed into the PBS. (15 minutes)
3. Cell dispensing: Load the cells field file. If multiple conditions/cell types are being dispensed, the user must manually edit file specify the location of each condition. This can be done by simply erasing spots where the cells are not to be dispensed, dispensing cells, and reloading the file and performing same edit for next condition. Load 15 *µl* into any well of the 384 well plate and manually aspirate 10 *µl* from the chosen well. To dispense cells use the CellenONE Basic method and adjust diameter and elongation parameters to gate for the relevant population. Incubate cells for 10 minutes post dispensing of last cell in 100 % DMSO for lysis. (*∼* 100 minutes for *∼* 2, 000 cells)
4. Evaporation Control: Several steps are taken to control evaporation. The humidifier is used to maintain humidity inside the chamber, and a cooling system is used to cool slides. Relative humidity inside the CellenONE is set to 75 % and the chiller temperature is set to dynamically chase one degree above the dew point. Next, load the perimeter field file. The SpotRun method is used to dispense system water in a perimeter surrounding each slide to provide further control for the local humidity of the reaction volumes. These humidity control settings are maintained throughout the entire next step (digestion) Re-dispense the perimeter throughout the 4 hour digestion as needed. (10 minutes)
5. Digestion: Load 30 *µl* into any two wells of the 384 well plate. Consecutively aspirate 20 *µl* from each well. Wash tip with the DipWash task. Load the Digest field file and dispense 12 nl of digestion buffer to each reaction volume using SpotRun. This brings reaction to a total volume of 20 nl. The digestion will proceed for 4 hours. (4 hours 20 minutes)
6. (Optional stop point) Let droplets dry out on slide, turn off humidity plate cooling to 10 degrees.
7. Labeling: Labeling protocols for TMT and mTRAQ reagents differ. Both protocols necessitate reconstituting labels in 100% DMSO. We do not store labels in DMSO long term, both TMT and mTRAQ are dried down using a speed vacuum on low heat and redissolved in 100% DMSO at the specified concentration. Both are used 1/3 diluted of the stock concentration. For TMT, the reaction can be performed in absence of aqueous buffer. If not starting from dried down droplets due to pause, turn off cooling and humidity before dispensing labels. Then load the label field file, load labels in wells G1-G14 and dispense labels with Pickup Sample and Dispense. For mTRAQ, leave or reset humidity and cooling settings as previously defined for digestion. Load the buffer field file and dispense 30 nl of 200 mM TEAB with SpotRun. Load the labels field file, load labels into wells G1-G3, and dispense labels with Pickup Sample and Dispense. Let labeling reaction proceed for 1 hour. (90 minutes)
8. Quenching: Pipette 30 *µl* of 5 % HA into any 2 wells of the 384 well plate. Consecutively aspirate 20 *µl* from each well. Load the quench field file and dispense 20 nL of 5 % hydroxylamine solution to each spot with SpotRun. Reset humidity and cooling controls are returned to previous settings as set during digest. After 20 minutes, another addition of 20 nL of 5 % HA.(50 minutes)
9. Prepare plate for sample collection: The final stage of sample prep is the automated collection of single cells into a 384 well plate for subsequent injection. The wells should optimally have a small volume in them to help facilitate the transfer of sample from the PDC to the well. If using carrier, pre-load 2 *µl* of carrier at concentration 1/2 desired concentration/*µl* into all wells J1-P24. If not using carrier or carrier is not yet ready, load 2 *µl* of water into all wells from J1-P24. In this study, the plate was pre-loaded with carrier when using TMT based workflow with isobaric carrier and pre-loaded with water for mTRAQ plexDIA workflow without used without carrier. The plate is then placed inside the cellenONE in the probe location on the right hand side. Humidity and cooling settings are maintained at 75 % relative humidity and plate cooling of 1 degree below dew point to prevent evaporation of liquid from wells during sample collection.
10. Sample pickup: First remove one of two nozzles. Sample clusters are pooled by using one PDC to sequentially aspirate droplets off the slide surface in 100 % acetonitrile solution via CellenONE PDC and syringe pump controls. Fill the wash tray with 100 % acetonitrile load the SamplePickup field file. If using higher multiplex as with the TMT labeling utilized in Scheme 1, use the Pickup off slide method which pools clusters of drops with 10 *µl*. For the mTRAQ labeling with only 3 cells per cluster as in scheme 2, we used the Pickup off slide smallvol method that pools samples with only 5 *µl*. Collecting one sample take approximately 50 seconds. This leads to a total pickup time of about 3.5 hours per 150 samples. (3.5 hours)
11. Storage/Sample injection: After pickup is complete, samples are completely dried down in the 384-well plate in a speed-vacuum on low heat. Then samples are either sealed with a foil cover and frozen at *−*80 *°C* for later analysis, or immediately reconstituted in a volume of 1.1 *µl* of either 0.1 % formic acid, or desired amount of carrier re-suspended in 0.1 % formic acid and sealed with a foil sheet. This 1.1 *µl* volume is used because we inject a volume 1 *µl* and the additional 0.1 *µl* prevents injecting air into lines. The plate is then placed in a compatible autosampler for mass spectrometry analysis. In our study, we utilized a Dionex 3000 which is compatible with 384 well plate based pickup. (20 minutes)

### Sample prep consumable cost estimate

The majority of consumable costs come from TMT labeling reagents. 10 *µl* of each label diluted 1/3 concentration from stock were aspirated for 2016 single cell droplets. This gives roughly 50 sample preparation amounts per batch of TMT. This brings the cost of TMT reagents per single cell to 8 cents. Glass slides can be purchased from scienion for 20 dollars per slide. About 500 single cells are prepared per slide, bringing the total cost to about $0.12*USD* per cell. Trypsin and other reagents used cost less than 1 cent per single cell.

### Preparation of cellenONE labeling validation samples

To estimate the labeling efficiency of for TMT dissolved in DMSO, we conducted two experiments. In the first experiment, we dispensed 50 ng of monocyte digest dissolved in HPLC grade water and 100 mM Triethyl ammonium bicarbonate buffer, pH 8.5. We also lysed, digested, and labeled 200 monocytes in 10 drops of 20 cells per drop following nPOP protocol. (Raw files AL078 and AL080 respectively)

### Mass spectrometry data acquisition

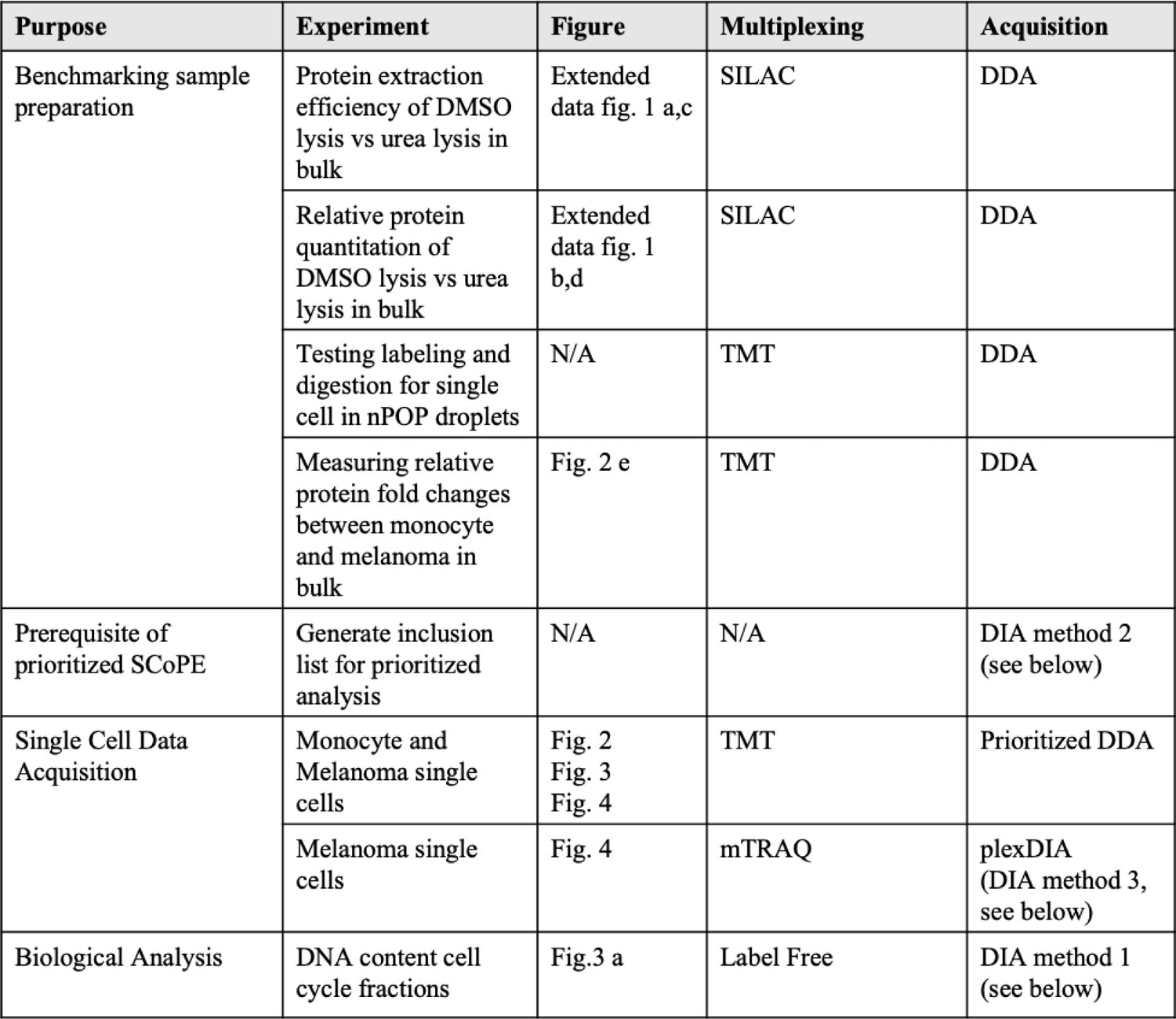

| Purpose | Experiment | Figure | Multiplexing | Acquisition |
| --- | --- | --- | --- | --- |
| Benchmarking sample preparation | Protein extraction efficiency of DMSO lysis vs urea lysis in bulk | Extended data fig. 1 a,c | SILAC | DDA |
|  | Relative protein quantitation of DMSO lysis vs urea lysis in bulk | Extended data fig. 1 b,d | SILAC | DDA |
|  | Testing labeling and digestion for single cell in nPOP droplets | N/A | TMT | DDA |
|  | Measuring relative protein fold changes between monocyte and melanoma in bulk | Fig. 2 e | TMT | DDA |
| Prerequisite of prioritized SCoPE | Generate inclusion list for prioritized analysis | N/A | N/A | DIA method 2 (see below) |
| Single Cell Data Acquisition | Monocyte and Melanoma single cells | Fig. 2<br>Fig. 3<br>Fig. 4 | TMT | Prioritized DDA |
|  | Melanoma single cells | Fig. 4 | mTRAQ | plexDIA (DIA method 3, see below) |
| Biological Analysis | DNA content cell cycle fractions | Fig.3 a | Label Free | DIA method 1 (see below) |

#### LC-MS analysis

All samples were were separated via online nLC on a Dionex UltiMate 3000 UHPLC using a 25cm x 75 *µl* IonOpticks Aurora Series UHPLC column (AUR2-25075C18A). For single-cell samples, 1 *µl* out of 1.1 *µl* of sample was picked up out of a 384 well plate (Thermo AB1384) placed on an auto sampler height adjuster for PCR plates (ThermoFisher 6820.4089) and sealed by an aluminum foil cover (ThermoFisher AB0626). Bulk samples were injected out of glass inserts (ThermoFisher 60180-509) sealed by rubber caps(ThermoFisher C5000-51B).

All samples except for plexDIA single-cells were run on a 95 minute total length method (53 minutes active gradients). 0.1 % formic acid in water was used for Buffer A and a 20% solution of 0.1% formic acid mixed in 80 % acetonitrile was used for buffer B. A constant flow rate of 200nl/min was used throughout sample loading and separation. Samples were loaded onto the column for 20 minutes at 1% B buffer, then ramped to 5 B buffer over two minutes. For unlabeled samples, the active gradient then ramped from 5% B buffer to 25% B buffer over 53 minutes. For labeled samples, the active gradient then ramped from 5% B buffer to 32% B buffer over 53 minutes. For all samples, gradient was then ramped to 95% B buffer over 2 minutes and stayed at that level for 3 minutes. The gradient then dropped to 1% B buffer over 0.1 minutes and stayed at that level for 4.9 minutes. plexDIA single cells followed the same gradient as other labeled samples, but ramped from from 5% B buffer to 32% B buffer over 25 minutes active gradient for a 60 minute total length method.

All samples were analyzed by a first generation Thermo Scientific Q-Exactive mass spectrometer. Electrospray voltage was set to 1.8 V, applied at the end of the analytical column. To reduce atmospheric background ions and enhance the peptide signal-to-noise ratio, an Active Background Ion Reduction Device (ABIRD, by ESI Source Solutons, LLC, Woburn MA, USA) was used at the nanospray interface. The temperature of ion transfer tube was 250 *°C* and the S-lens RF level was set to 80.

#### Standard DDA MS aquisition

After a precursor scan from 380 to 1600 m/z at 70,000 resolving power, the top 7 most intense precursor ions with charges 2 to 4 and above the AGC min threshold of 20,000 were isolated for MS2 analysis via a 0.7 Th isolation window with a 0.3 Th offset. These ions were accumulated for at most 300ms before being fragmented via HCD at a normalized collision energy of 33 eV (normalized to m/z 500, z=1). The fragments were analyzed by an MS2 scan with 70,000 resolution. Dynamic exclusion was used with a duration of 30 seconds with a mass tolerance of 10ppm.

#### DIA MS instrument methods

DIA instrument method 1: MS1 scans had the following parameters: 140k resolution, 3e6 AGC target, 512 maximum injection time, and a scan range from 450 to 1258Th. Two sets of DIA windows were used: 21 20Th-wide windows (spanning the space from 450Th to 860Th) and 8 50Th-wide windows (spanning the space from 859.5Th to 1256Th). The DIA windows had the following characteristics: 35k resolution, AGC target of 5e5, maximum injection time determined automatically, fixed first mass of 200Th, NCE of 33, and a default charge state of 2. DIA windows spanned the space from 450Th to 1256Th and included a 0.5Th overlap.

DIA instrument method 2: All characteristics of DIA Instrument Method 1 were preserved, but instead of two sets of variable-width DIA windows, three were used: 21 15Th-wide windows (spanning the space from 450Th to 755Th), 8 20Th-wide windows (spanning the space from 754.5Th to 911Th), and 7 50Th windows (spanning the space from 910.5Th to 1257.5Th). DIA windows spanned the space from 450Th to 1257.5Th and included a 0.5Th overlap.

DIA method 3: The duty cycle was comprised of one MS1 followed by three DIA MS2 windows of variable m/z length (specifically 120 Th, 120 Th, 200 Th and 580 Th) spanning 378–1,402 m/z. Each MS1 and MS2 scan was conducted at 70,000 resolving power, 3 × 10e6 AGC maximum and 300-ms maximum injection time.

#### Prioritized MS Data Acquisition Workflow

A prioritized analysis workflow[27] was used to increase consistency of identification and depth of coverage for the nPOP prepared single-cell data shown in Fig. 2, Fig. 3, and Fig. 4. A more in depth overview of steps in this workflow can be found in the prioritized analysis paper^27^. The steps towards creating the inclusion list are briefly outlined below.

1. A 10x concentrated injection of carrier and reference material was first analyzed via DIA instrument method 1 to generate a spectral library used to search subsequent 1x concentrated injections of carrier and reference material.
2. A 1x injection of carrier and reference sample was analyzed by DIA instrument method 2. The retention times and intensities from this run were used to create a prioritized inclusion list for subsequent prioritized single-cell analyses.
3. Three priority tiers were used, with the highest priority tier consisting of the most abundant peptides, and the lowest priority tier consisting of the least abundant. Additionally, a maximum of 4 peptides per protein were selected, choosing those 4 peptides with the highest precursor intensities, to maximize protein coverage.
4. This resulted in an inclusion list of 8,600 precursors. This list was then imported into MaxQuant.Live (v. 2.0.3 with priority tiers).
5. Several carrier and reference only samples were analyzed to further refine priority tiers of the inclusion list by reducing the list of 8,600 precursors base off peptides that were easily identifiable via prioritized data dependant acquisition analysis.
6. 2000 precursors were assigned to each of the three prioritized tiers, these precursors were then enabled for participation in the MaxQuant.live real-time-retention-time-alignment algorithm, and for MS2 upon detection. All remaining hard to identify or lowly abundant precursors that were not part of the priority tiers were then selected for participation in the MaxQuant.live real-time-retention-time-alignment algorithm, but not for MS2 upon detection.

### Interpreting raw mass spectra

#### Database searching of DDA and pSCoPE MS data

All raw data dependent acquisition/prioritized aquisition data were searched by MaxQuant^39, 40^ 1.6.17.0 against a protein sequence database including entries from the appropriate human SwissProt database (downloaded July 30, 2018) containing 20,386 proteins and known contaminants such as human keratins and common lab contaminants. For prioritized analysis, the fasta was limited to proteins with peptides featured on the prioritized inclusion list, `AndrewsnPOP_FASTA_v2.fasta` file. To make sure the limited fasta search was valid, we also searched a subset (20 MS runs) of the prioritized single cell data with a full fasta. Of the over 80,000 peptides confidently identified at 1% FDR with the full fasta search, only 15 peptides were not identified with the limited fasta search. This data is plotted in supplemental figure 8, S3. All other DDA runs were searched with a the full human fasta `swissprot_human_20211005.fasta`.

MaxQuant searches were performed using the standard work flow^41^. We specified trypsin specificity and allowed for up to two missed cleavages for peptides having from 7 to 26 amino acids. Methionine oxidation (+15.99492 Da) and protein N-terminal acetylation (+42.01056 Da) were set as a variable modifications. Carbamidomethylation was disabled as a fixed modification as cistine bonds were not carbamidomethylated. SILAC runs were searched in standard setting with multiplicity of 2. The heavy label was specified with Arg6 and Lys8. TMT labeled runs were searched Reporter ion MS2 with TMT labels specified in the `mqpar.xml`. TMT labeling efficiency searches were performed with TMTpro labels as variable modifications on lysine and n-termini. All peptide-spectrum-matches (PSMs) and peptides found by MaxQuant were exported in the msms.txt and the evidence.txt files. Lastly, we utilized Dart-ID to upgrade the confidence of PSMs that were consistently identified at the same retention time^42^. Results were filtered at 1% FDR by the dart qval parameter.

#### Analysis of Data Independent Acquisition MS data

The carrier and reference samples used to estimate retention times for the inclusion list of the prioritized analysis were then searched via Spectronaut’s (v. 15.4). The 10x concentrated carrier and reference sample was searched with directDIA feature, and TMTpro as a fixed modification on lysine residues and n-termini. *Note: TMTpro was not used for quantification or multiplexing in these DIA experiments, just for estimating the retention time for TMTpro labaled peptides.* The reference FASTA used for this analysis was `swissprot_human_20211005.fasta`. Default search parameters were used, with the following exceptions: source specific iRT calibration was enabled; Biognosys iRT kit was set to false; profiling strategy was set to template correlation profiling; Methionine oxidation (+ 15.99492 Da) was specified as a variable modification and TMTPro (+ 304.2071 Da) was specified as a fixed modification for peptide n-termini and lysines. From this run, a spectral library, `wAL00186.raw.kit` was generated. This spectral library was used to search 1x runs for generating accurate retention times for prioritized analysis of single cell sets.

Label free DIA runs of cell cycle fractions were searched with DIA-NN v1.8.0^43^ using an in silico fasta generated library enabled by deep learning. Raw files were searched together with match between runs (MBR). DIA-NN search settings: Library Generation was set to “IDs, RT, & IM Profiling”, Quantification Strategy was set to “Peak height”, scan window = 1, Mass accuracy = 10 ppm, and MS1 accuracy = 5 ppm, “Remove likely interferences”, “Use isotopologues”, and “MBR” were enabled. plexDIA single cell runs were searched using the plexDIA module in DIA-NN and a spectral library used to search single cells as described by Derks *et al.*^24^. Briefly, this library was generated by first running a bulk injection of 1,000 cell labeled with mTRAQ delta 8. This run was then searched using an in silico fasta generated library enabled by deep learning, with the mass of mTRAQ delta 8 appended to all lysine and n-termini in silico. The results of this search were used to generate a spectral library with roughly 5,000 human proteins.

### Preliminary downstream data analysis

#### SILAC data analysis of lysis validation data

When comparing relative protein levels in Jurkat and U-937 cells, SILAC ratios for peptides were computed by taking dividing each channel by its median, and then taking the ratio of the light and heavy channels, Extended Data Fig. 3. When comparing absolute abundances between heavy and light U-937 cells to measure efficiency of extraction, label swap experiments were ran so that both lysis conditions were measured with both heavy and light labels. The raw intensities for corresponding lysis methods were averaged and the ratio between different lysis methods was plotted, Extended Data Fig. 3.

#### Estimating labeling efficiency for TMT dissolved in DMSO

For labeling efficiency experiments (Raw files AL078 and AL080 respectively), we computed the labeling efficiency as the ratio of number possible lysine or n-termini that could be labeled containing TMTpro labels divided by number of possible labeling sites.

#### Assessing pSCoPE single cell digestion relative to the carrier

When using an isobaric carrier for single cell proteomic analysis, the missed cleavage rates of identified peptides likely reflect the missed cleavage rate of the carrier used for analysis. This makes assessing digestion quality of single cells in this context less straightforward. To get around this issue, we took the ratio of raw reporter ion intensity for all peptides identified that had corresponding missed cleaved versions also identified. These ratios in single cells were then compared to the same ratio from the carrier. If these two ratios are similar, it indicates that the the single cells are similarly well digested to the carrier.

#### Single-cell filtering and normalization of pSCoPE data

The single-cell data were processed and normalized by the SCoPE2 pipeline^14, 25^. This pipeline is also implemented by the scp Bioconductor package^44, 45^. Briefly, single cells with suboptimal quantification were removed prior to data normalization and analysis based on objective criteria: The internal consistency of protein quantification for each single cell was evaluated by calculating the coefficient of variation (CV) for proteins (leading razor proteins) identified with over 5 peptides for that cell. The coefficient of variation is defined as the standard deviation divided by the mean. The CVs were computed for the relative reporter ion intensities, i.e., the RI reporter ion intensities of each peptide were divided by their mean resulting in a vector of fold changes relative to the mean. Cells that fell outside the distribution were removed from analysis with a threshold of 0.41. Data was normalized as by procedure outlined by Specht *et al.*^14, 46^.

#### Single-cell filtering and normalization of plexDIA data

The report.pr matrix channels ms1 extracted output file from DIA-NN was used for quantitation. The single cell data was first filtered to 1% protein FDR with the Lib.PG.Q.value column of the DIA-NN output, and 1% Channel.Q.value.

#### Principal component analysis for single cell data sets

For Fig. 2e,f, Extended Data Fig. 9a, Extended Data Fig. 7a,c,d, and Extended Data Fig. 8a,b, we performed weighted principal component analysis in the space of all quantified proteins. Missing values were imputed using k-nearest neighbors imputation with k = 3. From the protein x single cell matrix, all pairwise protein correlations (Pearson) were computed. Thus, for each protein, there was a computed vector of correlations with a length the same as the number of rows in the matrix (number of proteins). The dot product of this vector with itself was used to weight each protein prior to principal component analysis. For figure all other PCA plots, principal component analysis was performed on the original data matrix without weighting.

#### Integration of pSCoPE and plexDIA melanoma datasets

To project plexDIA and pSCoPE data of melanoma cells into a joint space, each data set was first normalized to relative protein levels within the space of melanoma cells. Then, each data set was filtered for the 350 proteins that overlapped between data sets. Lastly, data was integrated using ComBat^47^.

### Cell cycle analysis

#### Bulk DNA sorting for CDC fraction sample preparation

Melanoma and monocyte cells were incubated using Vybrand Dye Cycle (Thermo V35003) following manufacturers instructions to stain for DNA content. Cells were sorted via the Beckman CytoFLEX SRT based off peaks corresponding to G1, S, or G2 phase of the cell cycle, Fig. 3a. Because the cells were stained with a membrane permeable dye, cell membranes stayed largly intact during sorting. This allowed us to pellet cells post sorting, remove PBS/sorting fluid, and resuspended in water at a concentration of 2,000 cells/*µl*. Cells were then frozen at −80 degrees Celsius for 10 minutes and then heated to 90 C for 10 minutes for lysis^18^. Proteins were then digested over night in a solution of 15 ng/*µl* of trypsin, 100 mM TEAB, and 1x Benzonase. Samples were then dried down and re-suspended in 0.1 % formic acid at a concentration of 2,000 cells worth of digest per *µl* for LC-MS analysis.

#### Identifying differential proteins from bulk CDC fractions

Differential protein abundance testing was performed using precursor-level quantitation. To account for variation in sample loading amounts, precursors from each sample were normalized to their sample-median. Then, each precursor was normalized by its mean across samples to convert it to relative levels within cell type. The normalized relative precursor intensities from different phases from both mononocytes and melanoma were grouped. Peptides from individual protein groups and compared by ANOVA between cell cycle phases to estimate the significance of differential protein abundance. To correct for multiple hypotheses testing, we used the Benjamini-Hochberg (BH) method to estimate q-values for differential abundance of proteins.

#### Constructing CDC phase markers

Phase markers were constructed from proteins identify with differential abundant each CDC phase in both monocyte and melanoma cells. These proteins were first identified on the bulk level. To further narrow the list of proteins used to create phase markers, we used proteins that contained multiple, positively correlated peptides in the single cell samples.

Phase markers were then constructed by averaging the abundances of all possible combinations of observed values for 2 or 3 proteins corresponding to each phase of the cell cycle. We selected groups two markers for each CDC phase that were positively correlated. This served as validation as we expect proteins that are highly abundant in same phase will positively covary.

Markers were first constructed in the space of monocyte cells and correlations between markers were validated in melanoma cells Fig. 3 b. Having validated the protein markers, we averaged protein markers within phase for downstream analysis.

#### Identifying proteins that covary with CDC markers

To identify proteins that covary with the phase marker vectors, we correlated the phase marker vectors to the measured protein levels of each protein using Spearman correlation. We adjusted the distribution of p-values obtained from the Spearman correlation test using the BH method and filtered results at FDR *<* 1%.

#### Cell cycle PCA

Single cell data matrix without imputation was first filtered for CDC marker proteins used for analysis (Supp. File 5). Monocyte and melanoma cells were then normalized to mean relative protein levels within cell type. To compute the PCA, a cell X cell correlation matrix was first computed using only pairwise observations. We then took the first two eigenvectors of this matrix to plot as PC1 and PC2. Cells were then colored by the average abundance of proteins from each cell cycle phase in Fig. 3c.

#### Cell cycle protein set enrichment analysis

To identify functionally coherent sets of proteins that covary with the CDC phase markers, we correlated proteins within a gene ontology set to the constructed phase markers. For this analysis, we only correlated CDC phase markers to pairwise observed values. The data was filtered so that only proteins with at least observations were utilized. This resulted in 3 distributions of correlations for each cell type (one per CDC phase) and thus 6 distributions total for monocytes and melanoma cells. Each distribution contained all correlations from the proteins in a given GO term to a CDC marker for a phase (G1, S, or G2). These 6 distributions were then compared with ANOVA to calculate p-values similar to previously reported analysis^48^. Only GO Terms for which we had at least 4 proteins were analyzed. We then used the Benjamini-Hochberg method to estimate the corresponding q values (FDR; false discovery rare) for each GO term. Significant terms that had the highest, or lowest degree of similarity between cell types were then displayed in Fig. 3.

#### Assigning cells to CDC phase

We took a greedy approach to assign cells to a CDC phase. First, we created a vector comprised of length 3X the number of cells, where each value was the average abundance of the observed values for G1, S, or G2 marker proteins in a given cell. We then sorted this vector from highest to lowest correlation. We then iterated down the list and sorted cells into the G1, S, or G2 bin based off the phase of each value. We sorted 50 % of cells into the G1 bin, 25 % of cells into the S and G2 bins based off the distribution observed from the bulk FACS CDC sorting.

### Melanoma sub population analysis

#### Melanoma sub population protein set enrichment analysis

Protein set enrichment analysis was performed by comparing the distributions of abundances between protein groups defined by GO terms. Relative levels for these proteins were compared for cells assigned to Cluster A and B. The t-test was performed using only observed values in the data (no imputation). In order to test a given gene set, it was required that a given gene set had at least 4 proteins measured in the single cells and that the data completeness for each population exceeded 80 %. The distribution of p-values for all GO terms tested was corrected for multiple hypothesis testing with the BH method. Only GO terms were reported with q values less than 1 % were reported.

#### Differential protein abundance testing between melanoma clusters

Differential protein testing was performed by Kruskal–Wallis test on relative protein levels for each individual protein between cells expressing the protein in cluster A vs cluster B. The Kruskal–Wallis test was performed on only observed values in the data (no imputation). It was required that a given protein had at least observed measurements in least 50 % of cells. The distribution of p-values for each protein was corrected for multiple hypothesis testing with the BH method. Only proteins with Q value less than 1 % FDR were reported.

#### Glycogen assay

About 30,000 melanoma cells were sorted by FACS from each of the 2 subpopulations, clusters A and B. The population corresponding to cluster B was sorted by top 1% of CD49c abundance via red florescent antibody (Biolegend 343808). Cells corresponding to cluster A were sorted from 25-75% abundance range. The two sorted populations were then split evenly into 3 aliqots of 10,000 cells each. Glycogen content was measured by fluorometric assay (Sigma Aldrich MAK016-1KT) using the manufacture provided protocol.

#### Joint protein distributions and correlation matrix Fig. 5a,b

Single cell data was normalized relative to the mean within each protein and log2 transformed and imputed values were marked as NA. Abundances for the observed values were then plotted against each other. Pearson correlations were computed and reported, and p-values were determined using the spearman correlation test.

Single cell data were then filtered for cells only contained in population cluster A. We filtered the data for proteins that were found to be significantly differential between cluster A and cluster B that were involved in either glycolysis, or oxidative phosphorylation. This differential abundance testing was performed as described above. We then computed all pairwise pearson correlations and plotted the resulting matrix as a heatmap.

#### Gradient of cell states, Fig. 5c

We filtered the integrated data matrix of melanoma cells from pSCoPE and plexDIA for only proteins that were found to be significantly differential between cluster A and cluster B. This set of proteins can be found in Supp. file 9. We then performed PCA in the space of these proteins by taking the first two eigenvectors of the cell x cell correlation matrix. All proteins that were upregulated in cluster A (cluster A markers) were colapsed to the mean value. These values were then plotted against PC1 and used to color code the PCA. The same was done for markers of cluster B.

### Extended Data Figures

**Extended Data Fig. 1.**
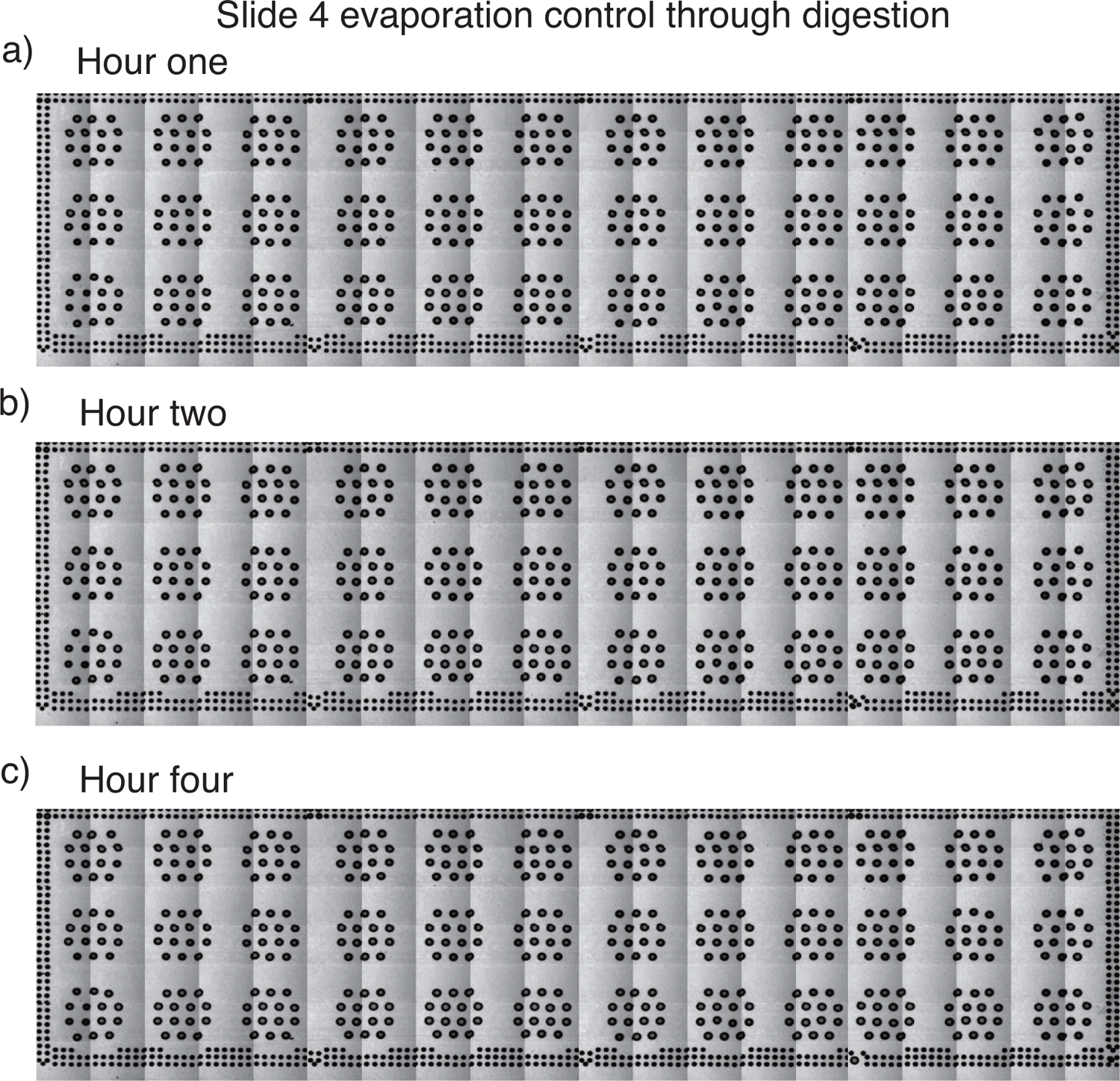
Evaluating droplet evaporation. Droplets on glass slide after adding DMSO, single cells, and digest mix after (**a**) one hour, (**b**) two hours, and (**c**) four hours.

**Extended Data Fig. 2.**
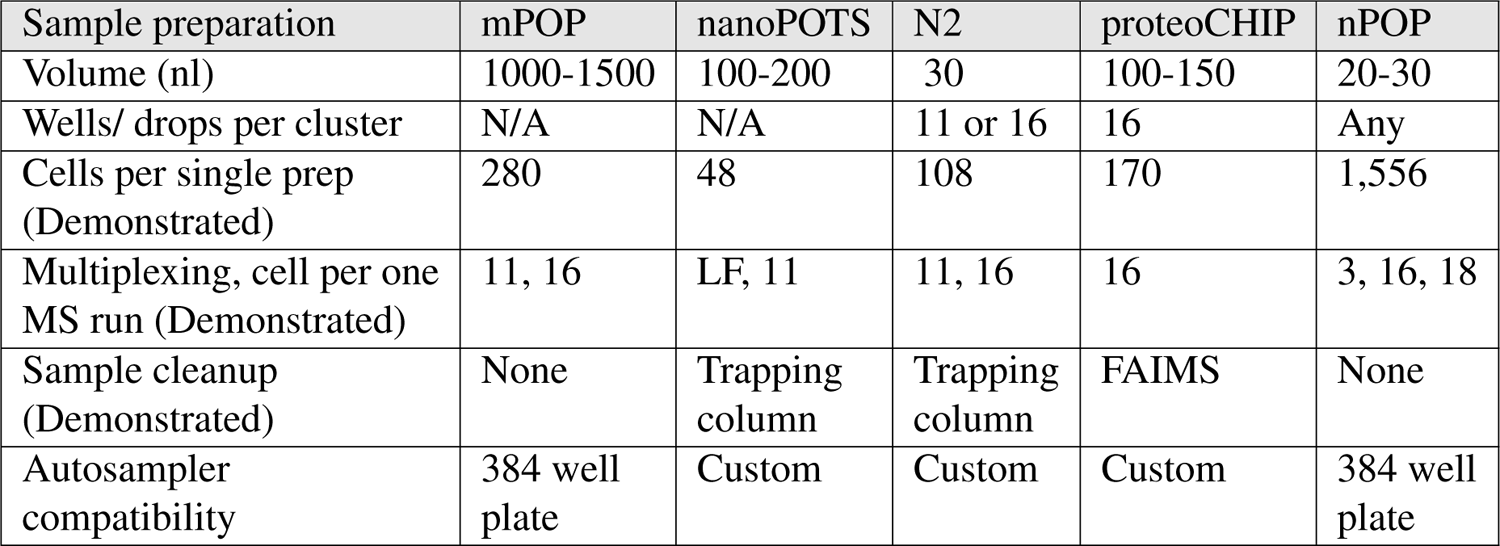
Table listing various important aspects of sample preparation for different single cell sample preparation methodologies.

**Extended Data Fig. 3.**
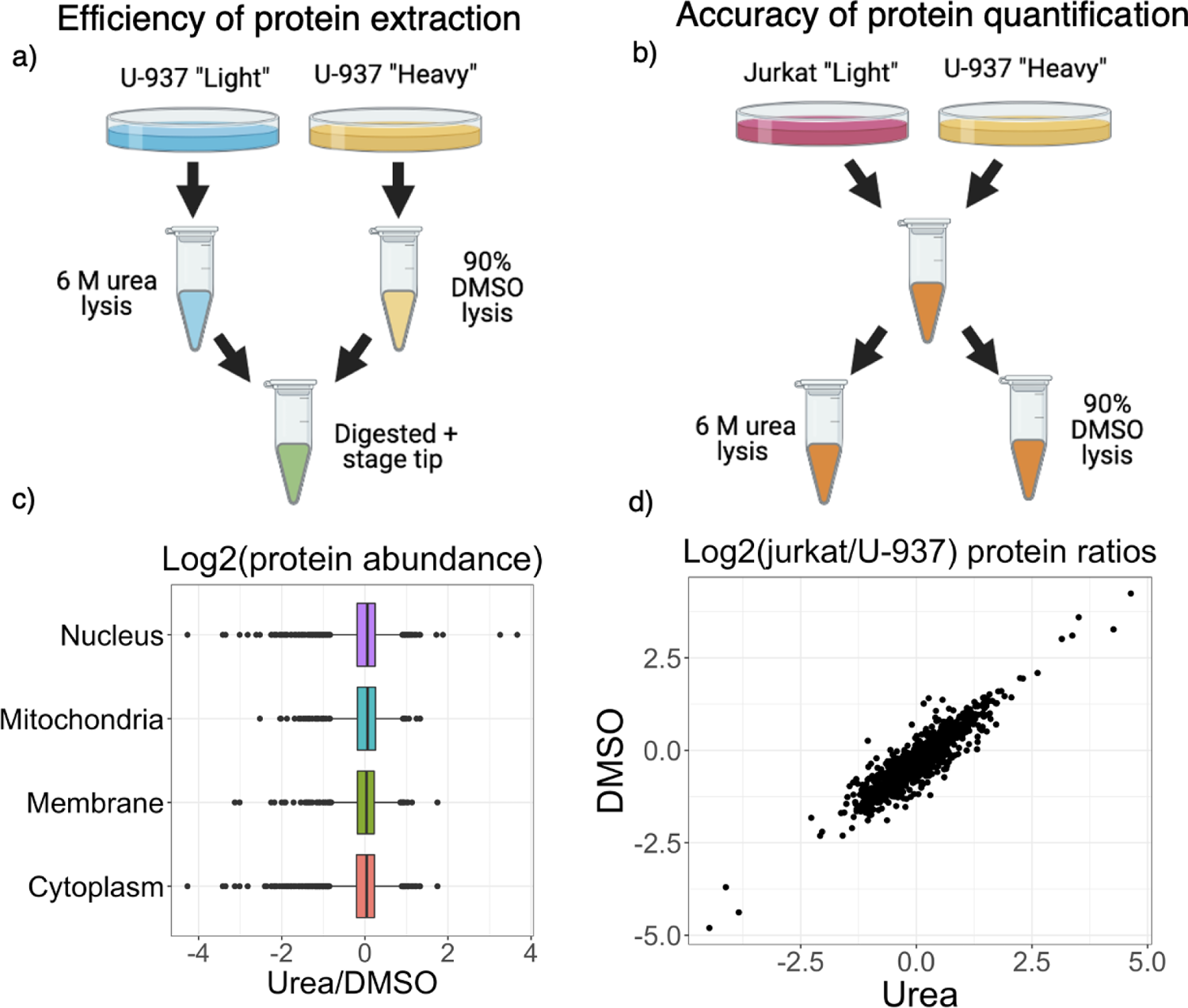
Evaluating the efficiency of protein extraction by DMSO cell lysis. (**a**) Equal number of U-937 cells labeled with “Light” and “Heavy” isotopes via SILAC were lysed with urea or DMSO, diluted, and combined for digestion. The SILAC ratios for proteins from different cellular compartments show comparable protein recovery for DMSO and urea cell lysis. (**b**) Equal number of SILAC labeled “Light” Jurkat and “Heavy” U-937 cells were combined, and the mixed sample was then divided for cell lysis either by urea or or by DMSO. The agreement between the SILAC ratios from the two methods supports the use of DMSO lysis for quantitative protein analysis.

**Extended Data Fig. 4.**
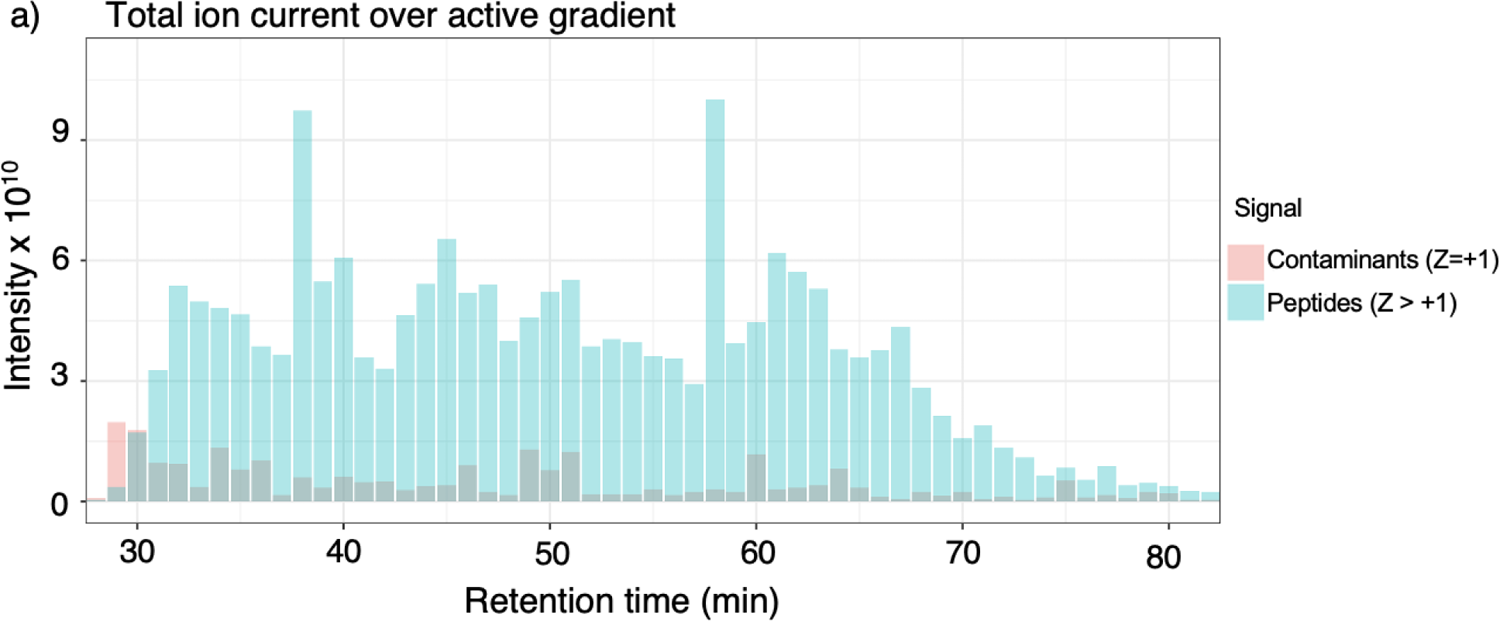
Single-cell samples processed by nPOP have low contamination and complete protein digestion. (**a**) Ion current from +1 charged ions (likely corresponding to contaminants) and *≥* +2 charged ions, likely correspond to peptides. Current from *≥* +2 charged ions far exceeds that of +1 indicating that contamination of samples prepared by nPOP is low.

**Extended Data Fig. 5.**
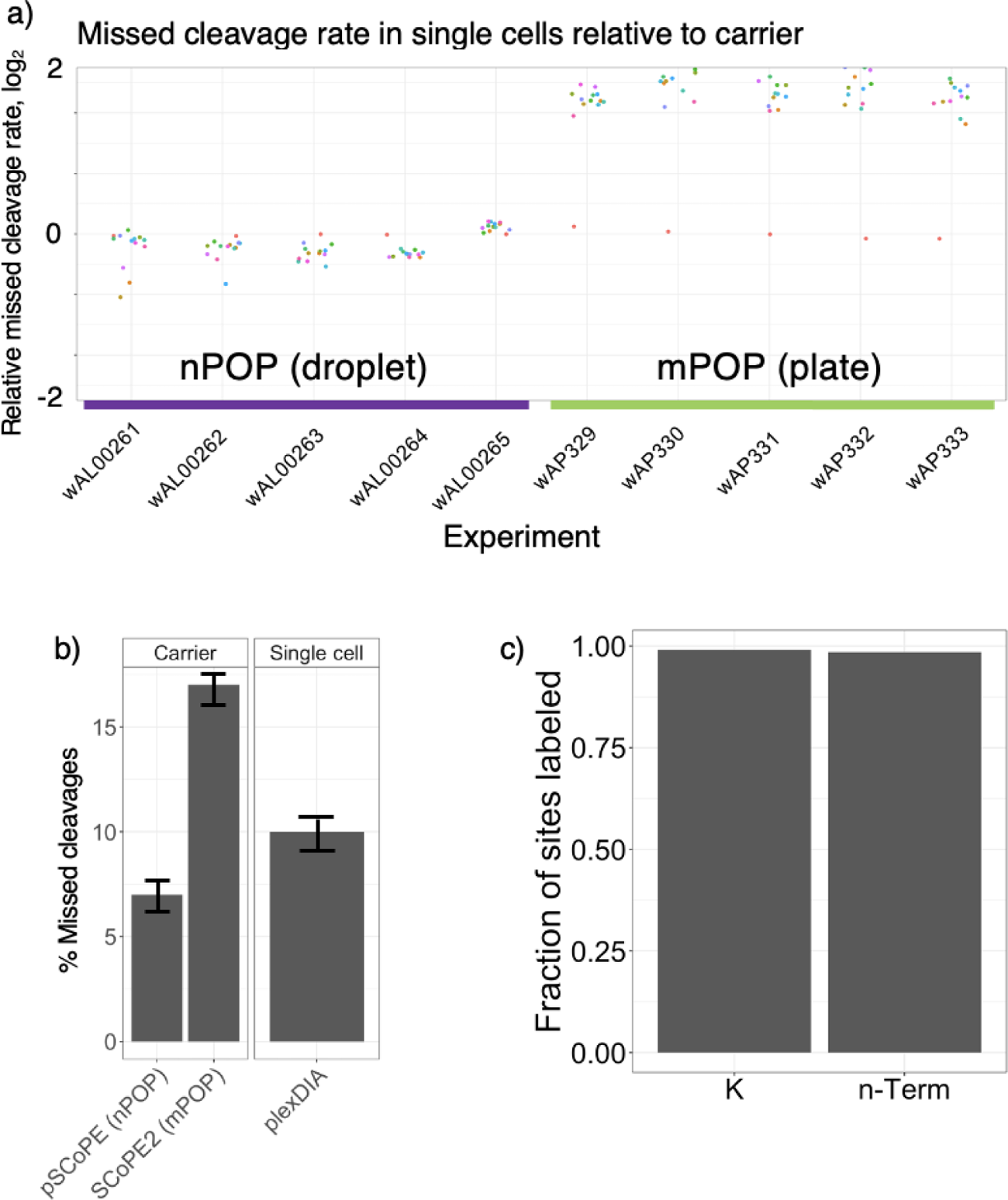
Single cell digestion evaluation. (**a**) The ratio of intensities for peptides with missed cleavages to their corresponding fully cleaved products is calculated in both the single cells and the carrier. The ratio of these two numbers is plotted as a way to evaluate miss cleavage rate in single cells compared to the carrier. Ratios centered around 0 indicate the proteins from single cells are digested as well as the proteins from the carrier sample, which has about 3 % peptides with missed cleavages. This panels is generated by DO-MS^49^ and can be found in the associated DO-MS report. Evaluating the metric in (**a**) depends on the digestion efficiency of the carrier. SCoPE2 data is taken from the data in the SCoPE2 protocol^25^. (**b**) shows that the carriers for the nPOP sets have a lower missed cleavage rate than the carrier for the mPOP sets increasing the significance of the better digestion in the single cells relative to the carrier. (**b**) also shows low missed cleavage rate for the single cells run without a carrier via DIA of *∼* 10%. (**c**) Fraction of labeled lysine or n-Termini sites out for many pooled single cells prepared by nPOP, labeled with TMTpro and searched with TMTpro as a variable modification.

**Extended Data Fig. 6.**
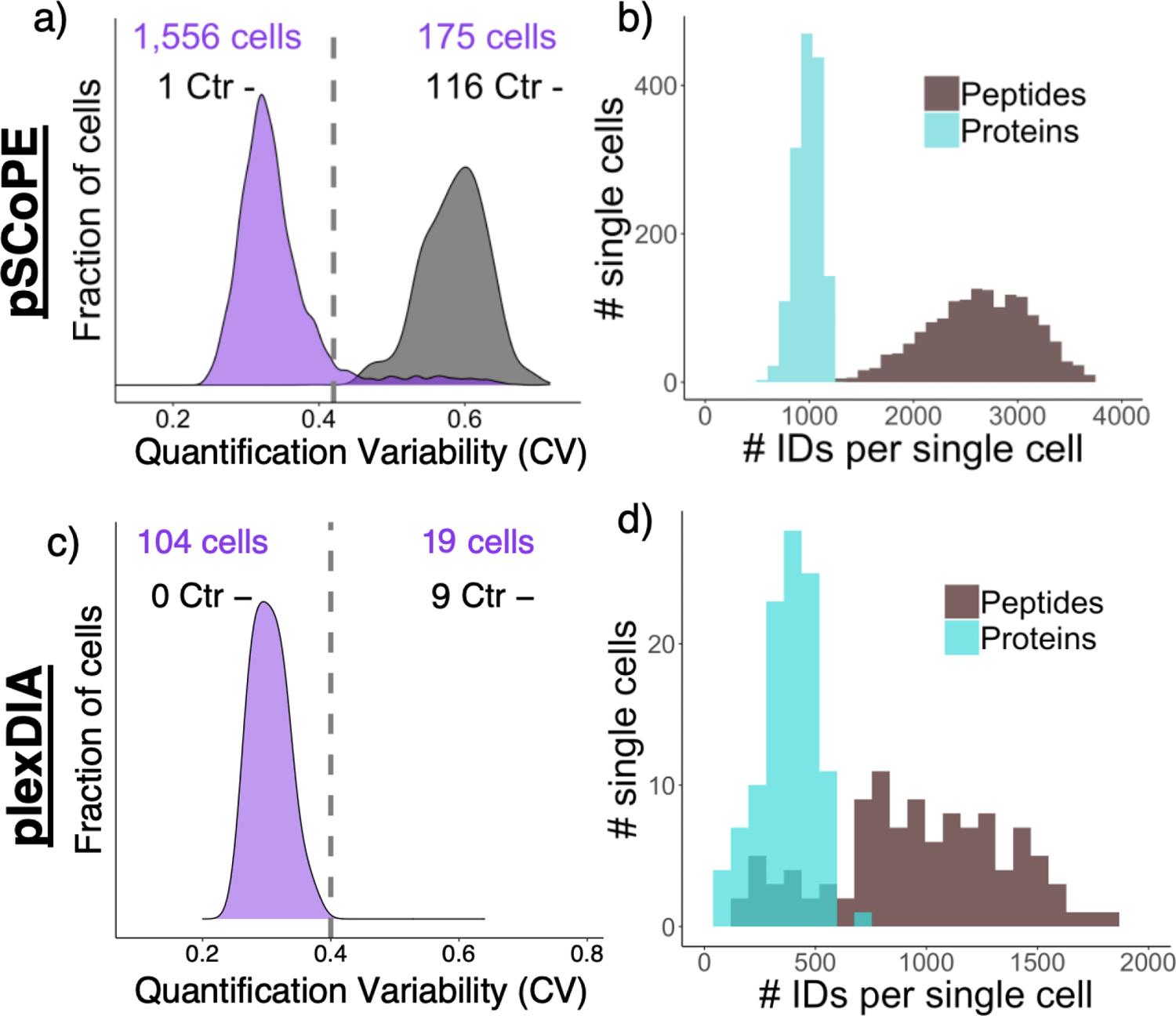
Single-cell data quality controls. (**a**) Quantitative variability for single cells analyzed by pSCoPE defined as the average CV per single cell between peptides mapping to the same protein. (**b**) Number of proteins and peptides per single cells analysed by pSCoPE. (**c**) Quantitative variability, CVs as described for (**a**), for single cells analyzed by plexDIA. (**d**) Number of proteins and peptides per single cells analysed by plexDIA.

**Extended Data Fig. 7.**
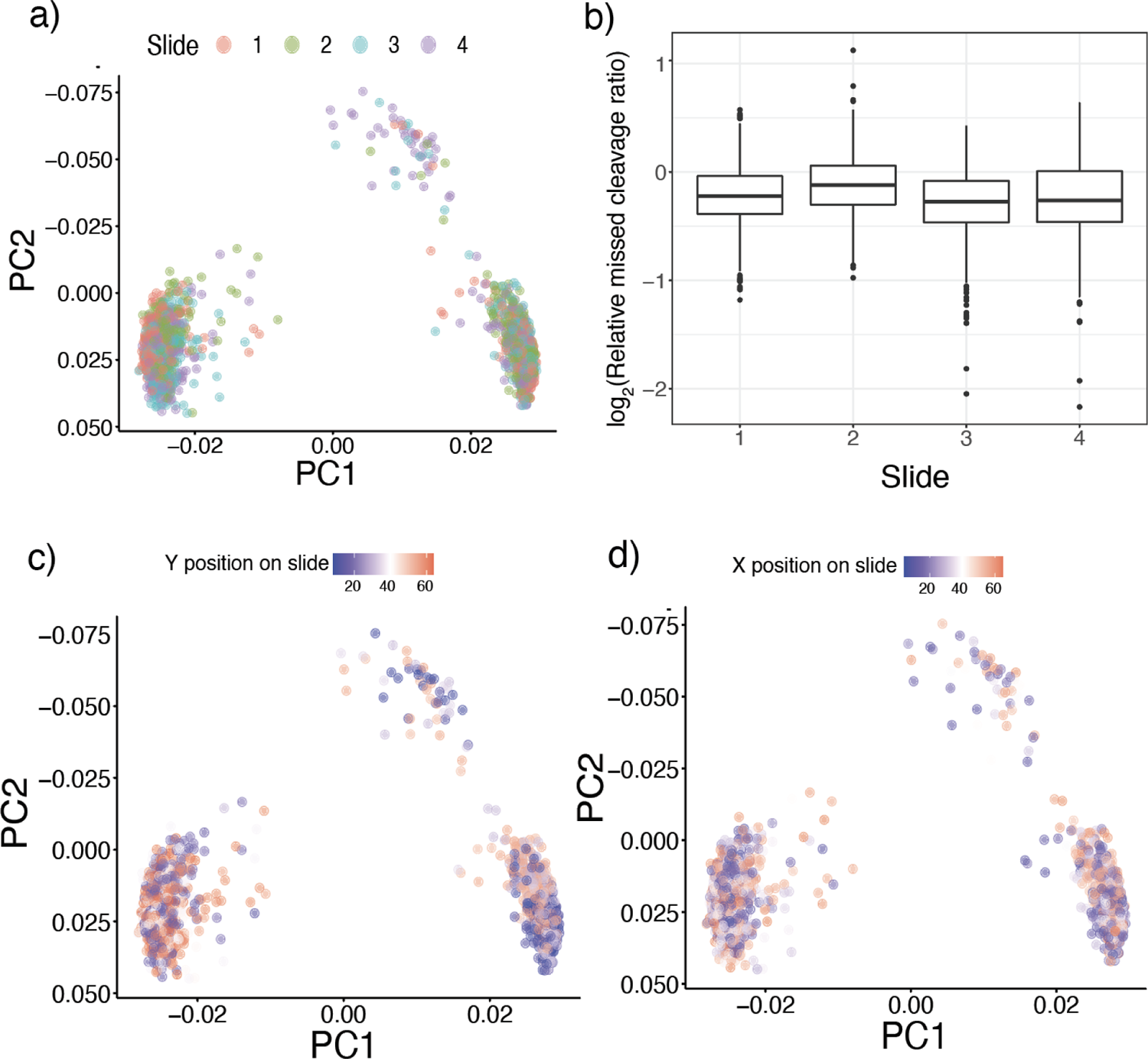
Examining potential batch effects. (**a**) PCA with single cells colored by which slide they were prepared on. (**b**) Distribution of ratios for missed cleaved peptides to their cleaved counterparts in single cells compared to the carrier sample plotted by slide. (**c**) Cells colored by X coordinates (left right) within slide. (**d**) cells colored by Y coordinates (front back) within slide.

**Extended Data Fig. 8.**
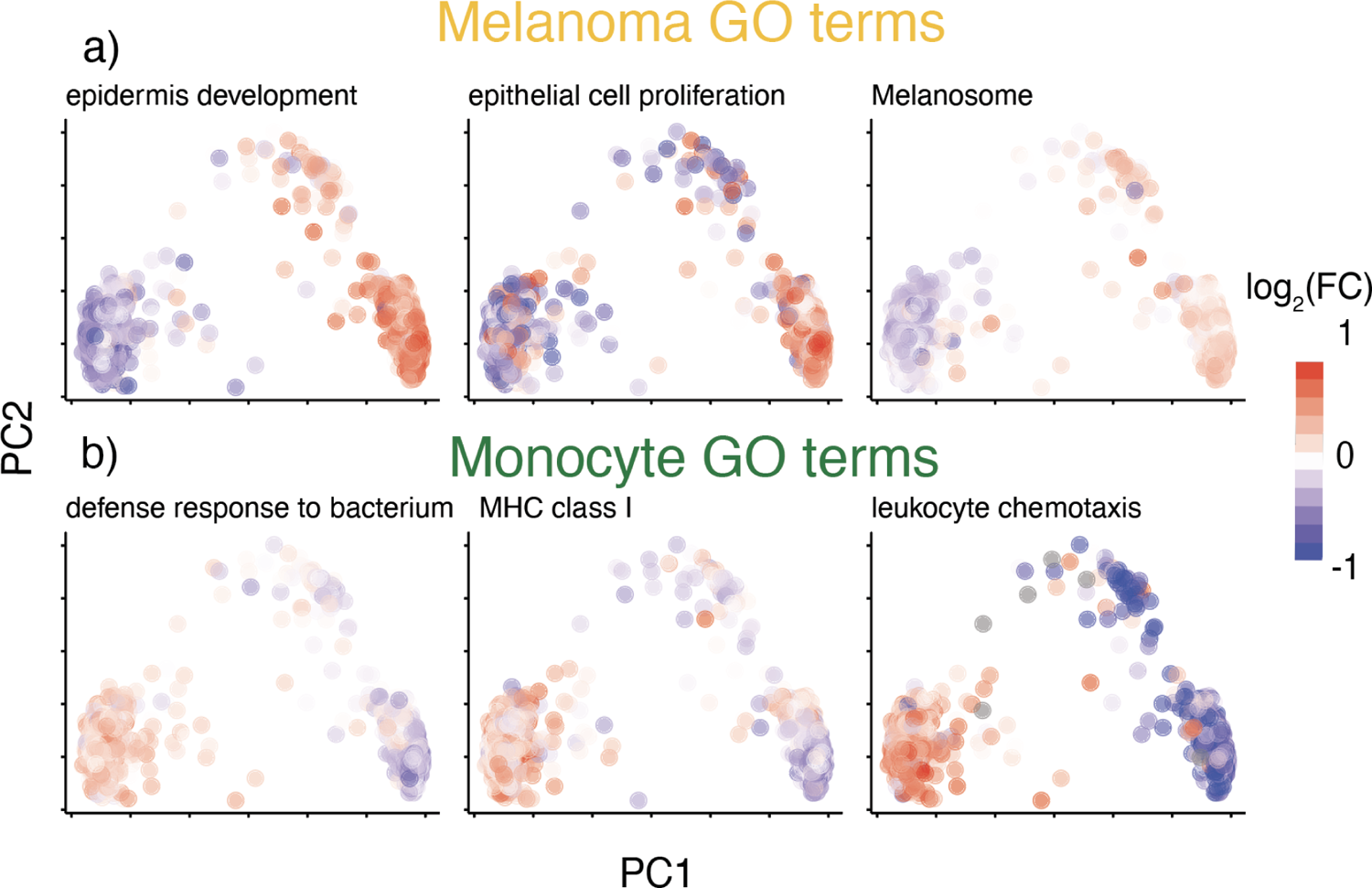
Protein set enrichment, Monocyte vs. Melanoma. Cells colored by the median abundance of proteins from protein sets associated with melanoma (**a**) and monocytes (**b**).

**Extended Data Fig. 9.**
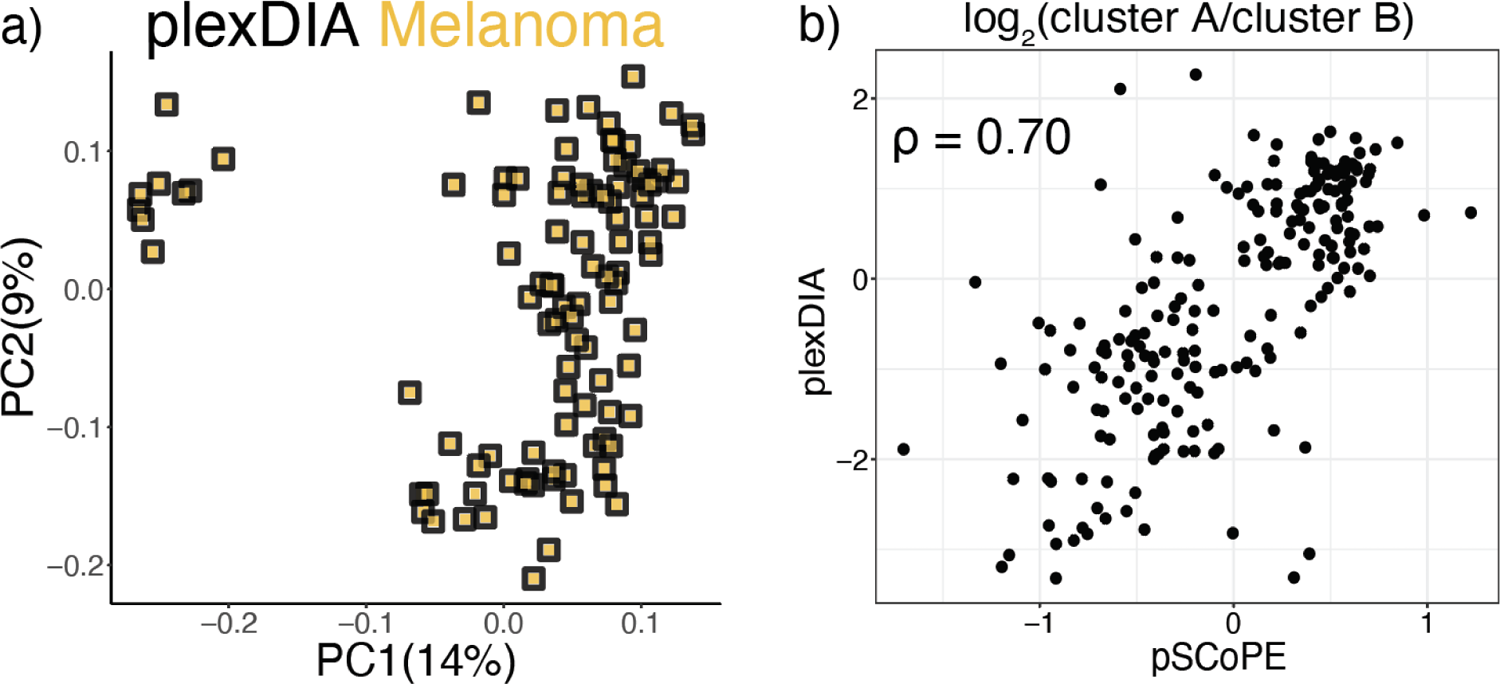
Validation of two clusters. (**a**) Principal component analysis of plexDIA melanoma cells showing two distinct clusters. (**b**) Protein fold changes between each cluster plotted in plexDIA vs pSCoPE data sets show agreement.

**Extended Data Fig. 10.**
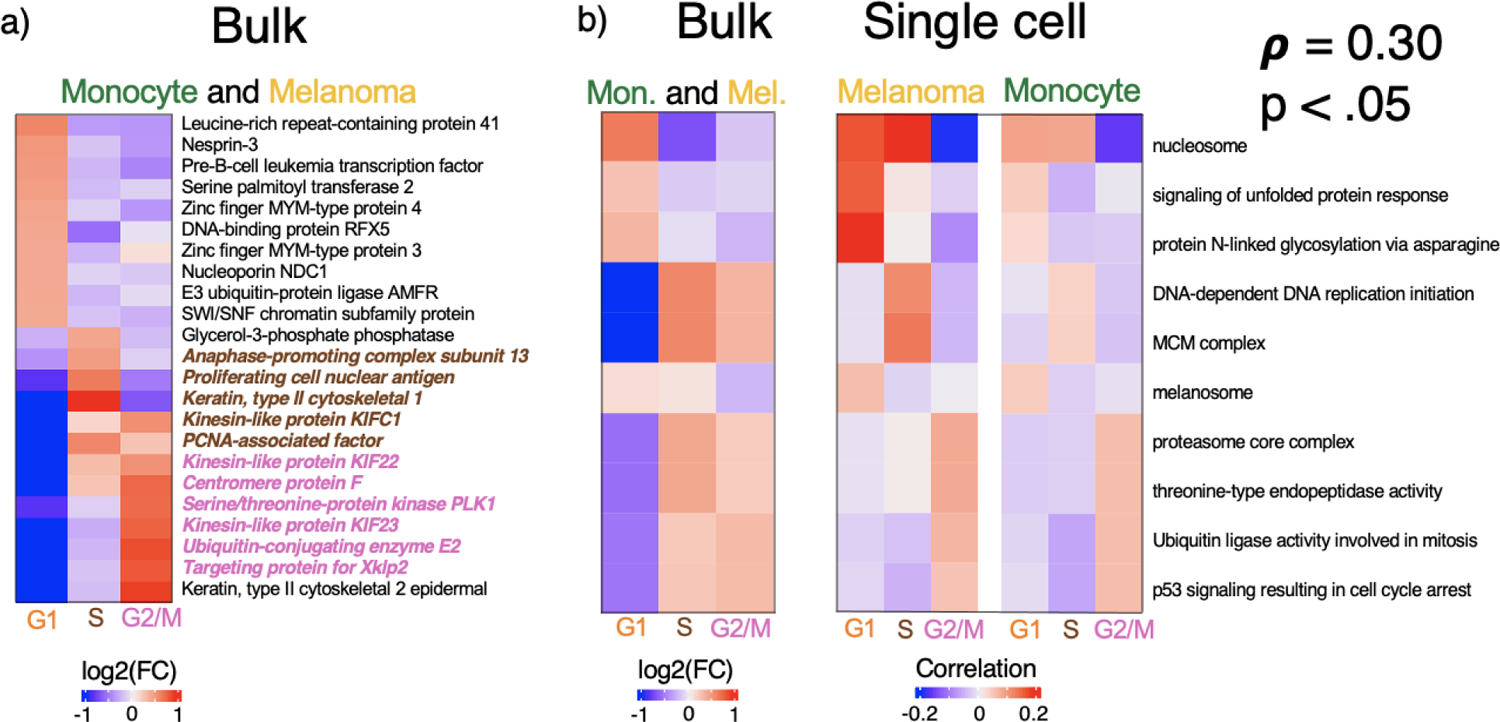
Cell cycle analysis quality controls. (**a**) Proteins found to be differential from both monocyte and melanoma bulk CDC fractions. Proteins previously reported to be involved in S phase are colored in brown and previously associated G2 proteins are colored in pink. (**b**) Protein set enrichment analysis for bulk data based off protein fold changes between three CDC fractions and corresponding significant protein sets from correlation based single cell protein set enrichment analysis. The two analysis are significantly correlated suggesting agreement between bulk and single cell analysis.

