## Supplementary material for "Exploring functional protein covariation across single cells using nPOP": Supplementary_data_5.html

DO-MS Report


### DO-MS Report

###### Version: 1.0.10 | PEP < 0.81 | Generated: 2022-08-18 13:37:35

#### Summary

##### MaxQuant Search Summary

MaxQuant Search Summary

MaxQuant Version: 1.6.17.0

Search Date: 04/08/2022 15:24:34

FASTA File: C:/Users/slavovlab/Desktop/AndrewsnPOP\_FASTA\_v2.fasta

##### Experiment Map

Map of raw file names to short names

##### MaxQuant Parameters

MaxQuant Search Parameters

##### MaxQuant Experiment Summary

MaxQuant experiment summary statistics

#### Chromatography

##### Elution profile: FWHM

Plotting the distrution of elution profile widths at half the maximum
intensity value for all observed ions.

##### Elution profile: FWHM of most abundant ions

Plotting the distrutions of elution profile widths at half the
maximum intensity value for the top ions with max intensity >
8e5.

##### Identification frequency across gradient

Plotting the frequency of peptide identification across the
chromatographic gradient.

##### Retention length of peptides at base

Plotting the retention length of identified peptide peaks at the
base.

#### Ion Sampling

##### Apex Offset

Plotting the distance from the peak of the elution profile the MS2
events were executed.

##### MS1 Intensity for identified ions

Plotting the MS1 intensity for all identified ions across runs.

##### MS1 Intensity for z>1 ions

Plotting the MS1 intensity for all peptide-like ions observed (not
necessarily sent to MS2) across runs.

##### MS1 Intensity for precursors in common

Plotting the MS1 intensity for precursors identified at any level of
confidence in all loaded experiments

##### MS1 intensity for identified peptides in common (normalized to reference)

Plotting the precursor intensity for peptides identified in all
experiments, normalized to the first experiment

##### MS1 Intensity for all ions

Plotting the MS1 intensity for all ions observed (not necessarily
sent to MS2).

##### Injection times, PSM resulting

Plotting distribution of MS2 injection times for scans that did
result in a PSM.

##### Number of MS2s per PSM

Plotting distribution of MS2 scans per PSM.

##### Injection times, no PSM resulting

Plotting distribution of MS2 injection times for scans that did not
result in a PSM.

##### MS1 fill-times (ms)

Plotting the distributions of MS1 fill times for scans with total ion
current greater than the 80th percentile of total ion currents.

##### MS2 scans per duty cycle

Plotting MS2 scans per duty cycle over the active gradient, defined
as retention time of first and last confidently IDd peptides (PEP <
0.01).

##### Identification rate faceted by duty cycle scan order and duty cycle length

Identification rate (peptides identified at PEP < 0.01 / all
scans) faceted by duty cycle scan order and duty cycle length

#### Peptide Identifications

##### Miscleavage rate, PEP < 0.01

Plotting frequency of miscleavages in confidently identified
peptides.

##### Miscleavage rate (percentage), PEP < 0.01

Miscleavage rate (percentage) for peptides identified with confidence
PEP < 0.01

##### Number of Confident Identifications

Plotting the number of peptides identified at each given confidence
level.

##### Precursor Ion Fraction (PIF)

The distribution of PIFs for identified peptides across all
experiments, where PIF is a measure of coisolation (1 = pure, 0 =
impure).

##### Fractional ID Rate

Fractional rate of confident PSMs and PSMs from all MS2 scan
events

##### Fractional ID Rate by intensity

Fractional ID rate by binned by MS1 intensity

##### Total ID Rate

Number of MSMS scans, PSMs, and confident PSMs

##### Error Probability Update

2D Densities of PSM error probabilities, given by MaxQuant (Spectra)
and DART-ID. Points below the 45 degree line indicate boosted confidence
(and lowered error probability), and vice versa for above the 45 degree
line. Set the PEP slider to 1 to see all PSMs regardless of initial
confidence.

```
## [1] "Plot failed to render. Reason: Error: Upload evidence_updated.txt\n"
```

##### Increase in Confident PSMs

Fold-change increase of PSMs at given confidence thresholds (in this
case, FDR thresholds)

```
## [1] "Plot failed to render. Reason: Error: Upload evidence_updated.txt\n"
```

##### RT Alignment Error

Alignment Error (Predicted RT - Observed RT) for the RT Alignment in
DART-ID

```
## [1] "Plot failed to render. Reason: Error: Upload evidence_updated.txt\n"
```

##### Peptide Identification Increase by Experiment

Increased number of peptide IDs per experiment, relative to existing
IDs before DART-ID

```
## [1] "Plot failed to render. Reason: Error: Upload evidence_updated.txt\n"
```

##### Fractional ID Rate

Fractional rate of confident PSMs and PSMs from all MS2 scan
events

```
## [1] "Plot failed to render. Reason: Error: Upload evidence_updated.txt\n"
```

##### Total ID Rate

Number of MSMS scans, PSMs, and confident PSMs

```
## [1] "Plot failed to render. Reason: Error: Upload evidence_updated.txt\n"
```

#### Contamination

##### Number of ions by charge state

Number of ions observed during MS1 scans by charge state

##### Summed precursor intensity by charge state

Plotting the summed precursor intensity by charge state.

##### MS1 Intensity, +1 ions

Plotting the intensity distribution of +1 ions.

##### m/z Distribution for +1 ions

Plotting the m/z distribution of +1 ions.

##### Intensity of z=1 across gradient

Plotting the intensity of z=1 ions observed.

##### Intensity of z=1 across gradient

Plotting the intensity of z=1 ions observed.

#### SCoPE-MS Diagnostics

##### Fraction of Missing Data per TMT Channel

This plot displays the fraction of missing values per reporter ion
per run per TMT channel, reported as 0 by MaxQuant.

##### Median intensity per TMT Channel, log10

This plot displays the intensity of reporter ion per run per TMT
channel.

##### rRI (misscleaved) / rRI (cleaved) in single cells

rRI defined as each channel divided by the carrier channel. Carrier
channel defined as channel with max median intensity. Plotting the ratio
of the mean relative reporter ion signal for miss-cleaved peptides / the
same for well-cleaved peptides

##### Mean reporter ion intensity for all channels but carrier

Plotting the mean reporter ion intensity for all channels but the
carrier. Carrier channel determined as channel with max median
intensity.

##### rRI (hydrophilic) / rRI (hydrophobic) in single cells

rRI defined as each channel divided by the carrier channel. Carrier
channel defined as channel with max median intensity. Plotting the ratio
of the mean relative reporter ion signal for hydrophilic peptides / the
same for hydrophobic peptides.

##### Spearman correlation of carrier channel to all other channels

Plotting spearman correlation of carrier channel to all other
channels. Carrier channel determined as channel with max median reporter
ion intensity.

##### Mean reporter ion intensity of synthetic peptide in all channels but carrier

Plotting the mean reporter ion intensity for spiked in peptides.
Carrier channel determined as channel with max median intensity.

```
## [1] "Plot failed to render. Reason: \033[1m\033[33mError\033[39m in \033[38;5;232m`dplyr::group_by()`\033[39m:\033[22m\n\033[38;5;232m\033[33m!\033[38;5;232m Must group by variables found in\n  `.data`.\nColumn `Raw.file` is not found.\nColumn `variable` is not found.\033[39m\n"
```

##### Single experiment only:

Reporter Ion Intensities vs. Carrier Intensities {.plot-title}

Comparing the reporter ion intensities for all TMT channels to the
carrier channel, chosen automatically as the most intense channel
(median intensity).

```
## [1] "Plot failed to render. Reason: Error: Please select a single experiment\n"
```

##### Single experiment only:

Reporter ion channel spearman correlations {.plot-title}

Plotting all pairwise TMT reporter ion intensity channel correlations
(Spearman) for a single run.

```
## [1] "Plot failed to render. Reason: Error: Please select a single experiment\n"
```

#### Carrier

##### TMT Labeling Efficiency, PEP < 0.01

Comparing relative rates of IDs for peptides with, or without the TMT
tag. Only compatible with searches performed with TMT as a variable mod
(n-terminus and on lysine)

```
## [1] "Plot failed to render. Reason: Error: Upload evidence.txt in labeling efficiency input\n"
```

##### Labeled / unlabeled precursor ion intensity, PEP < 0.01

Labeled / unlabeled precursor ion intensity. Only compatible with
searches performed with TMT as a variable mod.

```
## [1] "Plot failed to render. Reason: Error: Upload evidence.txt in labeling efficiency input\n"
```

##### Miscleavage rate percentage, PEP < 0.01

Percentages of missed cleavages for confidently identified peptides
(PEP < 0.01). Efficiency <20% desirable.

##### Miscleavage rate, R and K, PEP < 0.01

Miscleavage rate at arginine and lysine for peptides filtered to PEP
< 0.01
