## Supplementary figures and images for "Exploring functional protein covariation across single cells using nPOP"

### Supplemental_figure_8.png

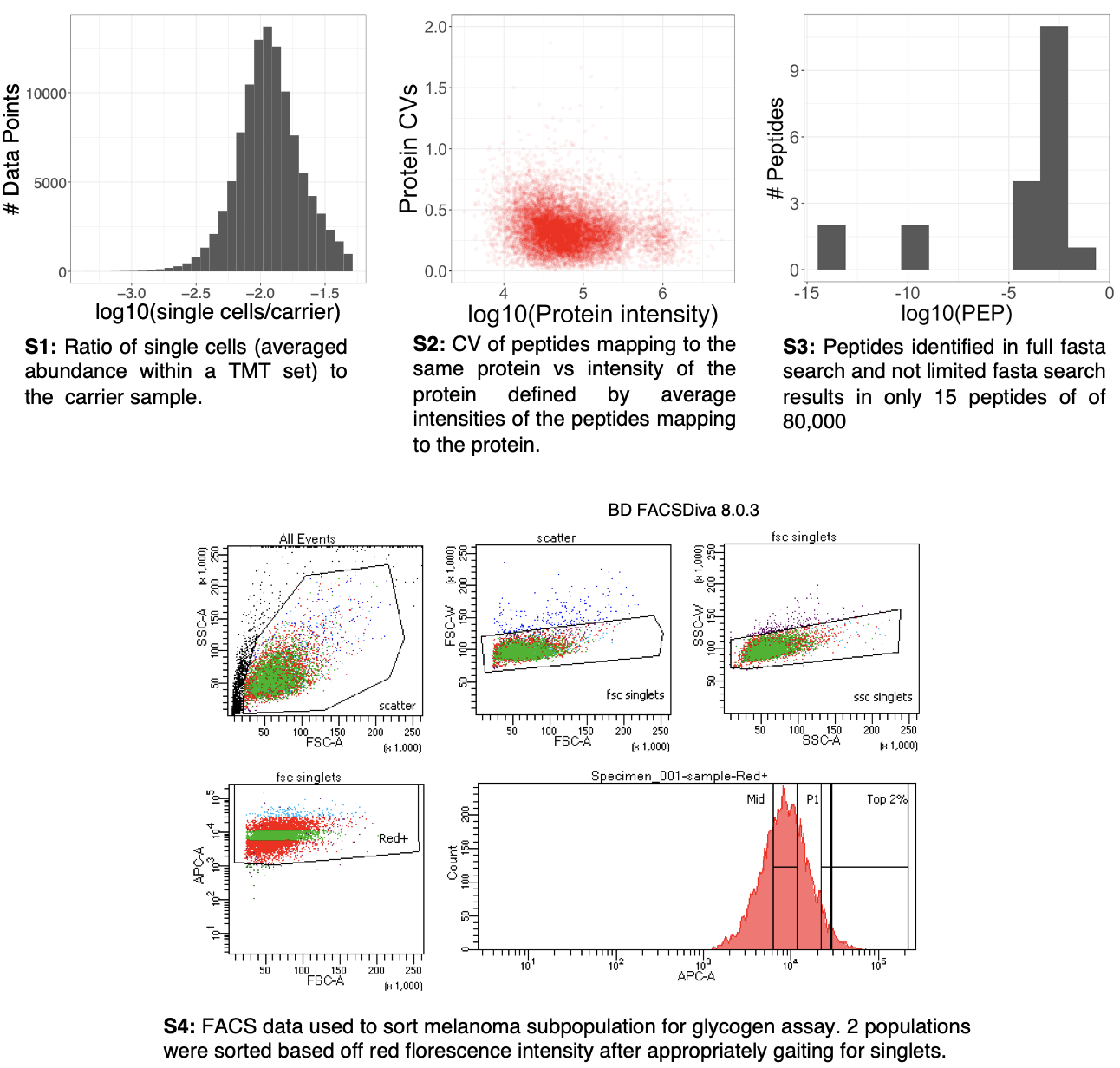
